# Using variation in arbuscular mycorrhizal fungi to drive the productivity of the food security crop cassava

**DOI:** 10.1101/830547

**Authors:** Isabel Ceballos, Ivan D. Mateus, Ricardo Peña, Diego Camilo Peña-Quemba, Chanz Robbins, Yuli M. Ordoñez, Pawel Rosikiewicz, Edward C. Rojas, Moses Thuita, Deusdedit Peter Mlay, Cargele Masso, Bernard Vanlauwe, Alia Rodriguez, Ian R. Sanders

## Abstract

The unprecedented challenge to feed the rapidly growing human population can only be achieved with major changes in how we combine technology with agronomy^1^. Despite their potential few beneficial microbes have truly been demonstrated to significantly increase productivity of globally important crops in real farming conditions^2,3^. The way microbes are employed has largely ignored the successes of crop breeding where naturally occurring intraspecific variation of plants has been used to increase yields. Doing this with microbes requires establishing a link between variation in the microbes and quantitative traits of crop growth along with a clear demonstration that intraspecific microbial variation can potentially lead to large differences in crop productivity in real farming conditions. Arbuscular mycorrhizal fungi (AMF), form symbioses with globally important crops and show great potential to improve crop yields^2^. Here we demonstrate the first link between patterns of genome-wide intraspecific AMF variation and productivity of the globally important food crop cassava. Cassava, one of the most important food security crops, feeds approximately 800 million people daily^4^. In subsequent field trials, inoculation with genetically different isolates of the AMF *Rhizophagus irregularis* altered cassava root productivity by up to 1.46-fold in conventional cultivation in Colombia. In independent field trials in Colombia, Kenya and Tanzania, clonal sibling progeny of homokaryon and dikaryon parental AMF enormously altered cassava root productivity by up to 3 kg per plant and up to a 3.69-fold productivity difference. Siblings were clonal and, thus, qualitatively genetically identical. Heterokaryon siblings can vary quantitatively but monokaryon siblings are identical. Very large among-AMF sibling effects were observed at each location although which sibling AMF was most effective depended strongly on location and cassava variety. We demonstrate the enormous potential of genetic, and possibly epigenetic variation, in AMF to greatly alter productivity of a globally important crop that should not be ignored. A microbial improvement program to accelerate crop yield increases over that possible by plant breeding or GMO technology alone is feasible. However, such a paradigm shift can only be realised if researchers address how plant genetics and local environments affect mycorrhizal responsiveness of crops to predict which fungal variant will be effective in a given location.

---

For millennia farmers have improved crops using naturally occurring intraspecific plant genetic variation to improve productivity. However, rates of yield increase attributed to plant breeding and GMO-crop technology are not considered sufficient to feed the projected global human population^1^. Beneficial soil microbes can potentially improve plant growth, but in sharp contrast to plant breeding, there has been little attempt to improve them. Innovative approaches to microbial management of symbiotic microorganisms could bring great benefits^5,6^. For decades, AMF have been known to increase plant growth^7^, although they are not consistently used in agriculture or the focus of an improvement program (Supplementary information note 1).

We investigated whether naturally occurring variation of a commonly occurring AMF species of agricultural soils (*Rhizophagus irregularis*) can significantly alter the productivity of the globally important food security crop cassava. Cassava feeds almost 800 million people in 105 countries and responds positively to AMF inoculation in farming conditions^4,8^.

Establishing a link between discernible patterns of genetic variation within a beneficial microbial species and plant growth has never been attempted, but if the link exists then patterns of microbial genetic variation can be used as a predictor for selection of beneficial strains. In AMF, mapping quantitative trait loci and genome-wide association studies are not currently realistic because of lack of recombinant lines, inadequate plant phenotypic datasets, and a limited panel of isolates. Another way to identify the existence of such a link is to demonstrate whether quantitative traits of a crop species are influenced by the genetic relationships among individuals of an AMF species, as discerned by phylogenetic relationships^9^. First, quantitative fungal growth traits were measured in a set of *R. irregularis* isolates, spanning the phylogeny of this species and using data on thousands of single nucleotide polymorphisms (SNPs) distributed widely across their genomes^10,11^. Second, we inoculated one cassava cultivar (NGA 16) with a set of 11 *R. irregularis* isolates also spanning the *R. irregularis* phylogeny. Indeed, the fungal isolates differed significantly in their quantitative growth traits (Supplementary information table S1) and there was a significant phylogenetic signal on spore density and clustering (Supplementary figure 1; Supplementary information table S1). The fungal isolates differentially colonized the roots and had a strong and significant differential effect on cassava growth, with the strongest effect on cassava root dry weight (Supplementary information table S1). Patterns of genome-wide variation among the isolates were significantly associated with root dry weight and total cassava dry weight (Figure 1; Supplementary information table S1). This establishes the first clear a link between genetic signatures of a beneficial microorganism and growth of a globally important crop plant.

**Figure 1.**
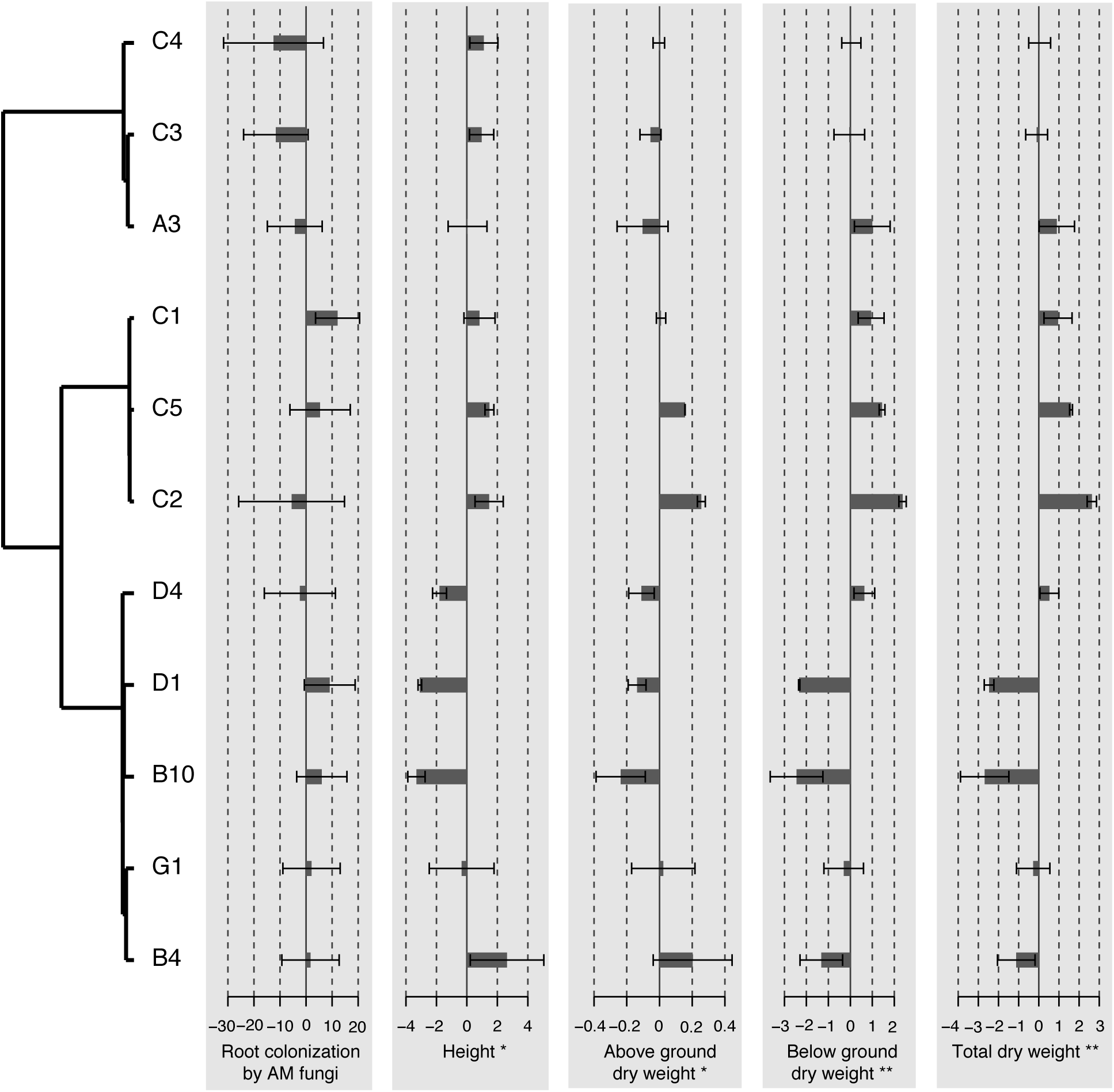
Variation in root colonization by AMF cassava height, shoot dry weight, root dry weight and total cassava dry weight among plants inoculated with 11 genetically different isolates of *R. irregularis* (experiment 2). Values shown are centred on the mean response across all treatments and bars represent the standard deviation. Value of traits for fungal inoculation treatments are arranged according to a dendrogram of genetic relatedness among fungal isolates. * *P* ≤ 0.05; ** *P* ≤ 0.01 represent significance of tests of a phylogenetic signal of a given trait with at least one test. Statistical tests for differences in a trait among treatments and tests for the existence of a significant phylogenetic signal are given in Supplementary information table S1.

Cassava has previously been shown to benefit from inoculation with *R. irregularis* under conventional cassava cultivation conditions in Colombia^8,12^ and represents a good candidate for an improvement program (Supplementary information note 1). While AMF isolates can affect plant growth in sterile soil, this is rarely validated in normal farming conditions. In a third experiment, we quantified how much variation in the growth of cassava (cultivar CM4574) occurred in normal farming conditions in Colombia using 5 genetically different *R. irregularis* isolates^10^. Cassava root weight ranged from 4.0 – 5.9 kg plant^-1^ among inoculated plants, with non-inoculated plants falling in the middle of this range (Figure 2; Supplementary information table S2 and note 2). This translates to a projected yield alteration of up to 18.6 tons ha^-1^ fresh root weight between treatments inoculated with genetically different isolates (Supplementary information note 3). Because isolates have been *in vitro* cultured for 16 years in an identical environment, results imply genetic variation (and possible additional epigenetic variation) in AMF leads to crop production differences (Supplementary information note 4).

**Figure 2.**
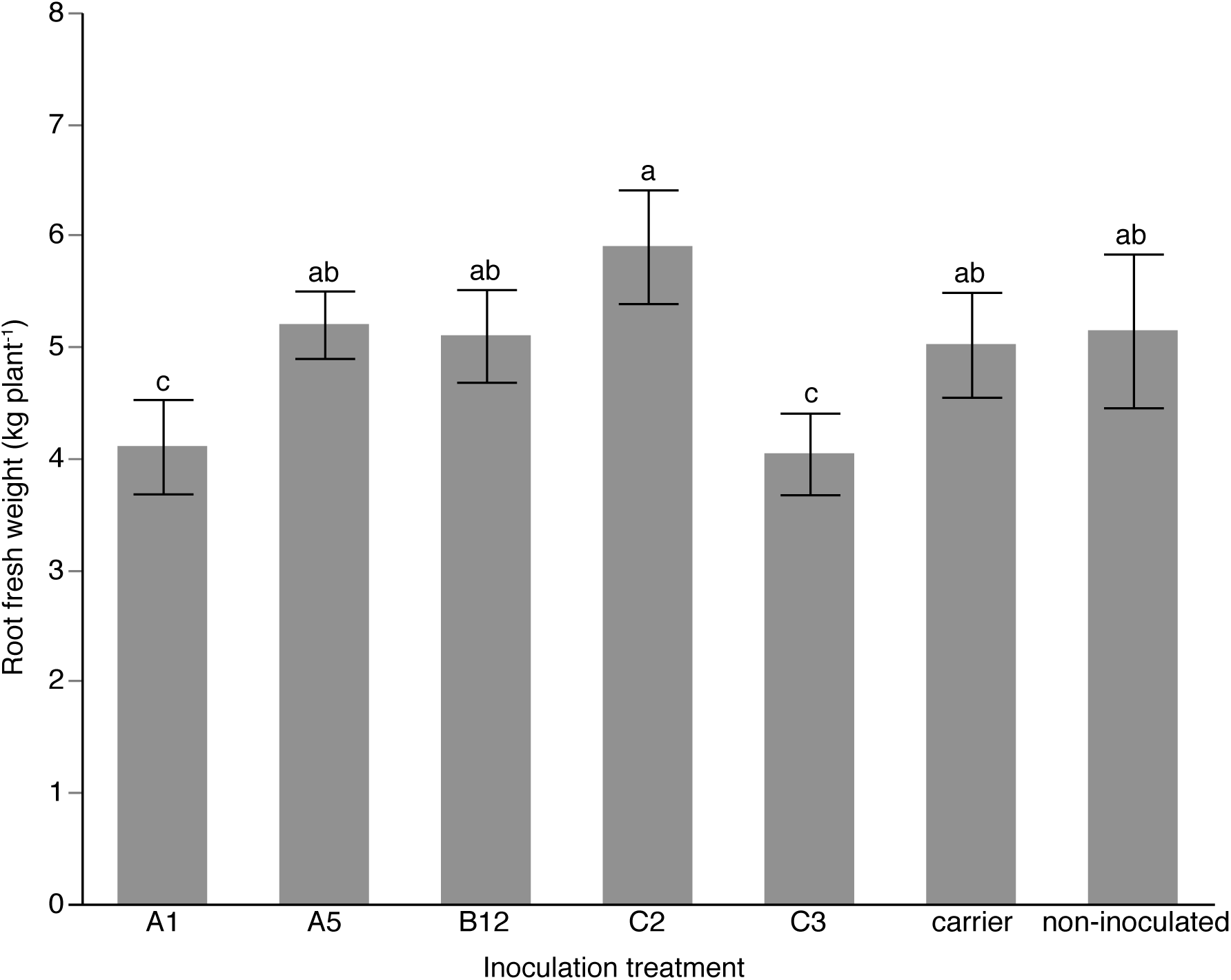
Mean root fresh weight per plant of cassava grown in normal farming conditions in Tauramena, Colombia and with 7 different inoculation treatments (experiment 3). Each value represents the mean of 6 replicate plots where the value o for each plot is the pooled mean of several plants. Bars represent ±1 S.E. Plants were inoculated with *R. irregularis* isolates A1, A5, B12, C2, C3 or were inoculated with the carrier without *R. irregularis* or were not inoculated. Different letters represent significant differences at *P* ≤ 0.1. See Supplementary information table S2 for statistical analyses.

Clonally produced single spore progeny of an *R. irregularis* isolate strongly altered the growth of rice in sterile conditions (Supplementary information note 5)^13^. A fourth experiment tested the effect of single spore progeny of two parental *R. irregularis* isolates on cassava root productivity in conventional cassava cultivation conditions in Colombia (Supplementary information note 5 and table S3). The same experiment was conducted in successive years, using two varieties of cassava. Cassava root fresh and dry weight was differentially affected by sibling fungi but how they responded differed between cassava varieties, with one variety (CM4574) responding more strongly to the fungal siblings (Supplementary figure 2; Supplementary information table S4). Results from the successive trials showed that the fungal treatments substantially affected cassava root weight under normal cassava cultivation in Colombia in a reproducible way over time, even when the non-inoculated control was removed from the analysis (Supplementary information table S4). The fundamental question of this experiment was to test whether variation among progeny of parental *R. irregularis* isolates significantly altered cassava root productivity and to measure the potential amplitude of the effect. Because the two parental isolates are genetically different from each other, analyses across all treatments is not appropriate to answer this question. Due to a strong AMF inoculation treatment by cassava variety interaction, we compared cassava root weight among plants inoculated with a given parental isolate and its progeny and by cassava variety separately. Root fresh and dry weight significantly differed by as much as 2.44-fold in cultivar CM4574 among plants inoculated with the parental isolate C2 and its progeny (Figure 3a-d; Supplementary information table S5). Inoculation benefit, a measure of the response to inoculation compared to non-inoculated plants, also differed significantly among the plants inoculated with progeny of C2 (Supplementary figure 3; Supplementary information table S5). Cassava inoculated with parental isolate C3 and its progeny showed a remarkable 2.94-fold difference in root weight, as well as differences in inoculation benefit in CM4574 (Figure 3e-h; Supplementary figure 3; Supplementary information table S5). AMF colonization of roots occurred in all treatments (Supplementary figure 2) and averaged over the two experiments, neither AMF treatments nor cassava cultivar significantly influenced mycorrhizal colonization (Supplementary information table S4). However, analysis of colonization in the 1^st^ trial revealed that colonization was significantly affected by inoculation treatments in cultivar CM4574 (data not shown).

**Figure 3.**
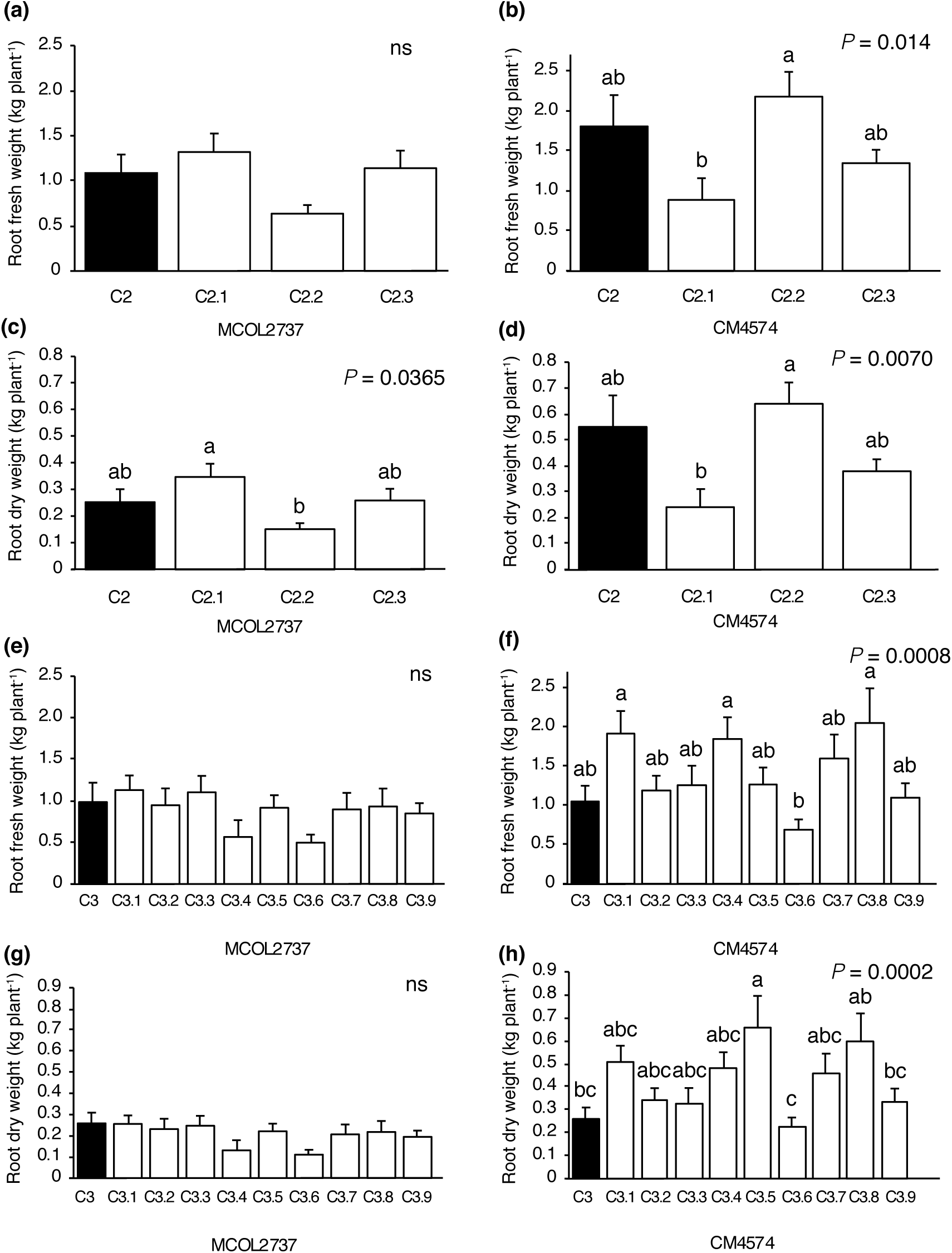
Mean root fresh and dry weight of cassava cultivars MCOL2737 and CM4574 inoculated with the parental *R. irregularis* isolate C2 and its clonal progeny and the parental *R. irregularis* isolate C3 and its clonal progeny averaged over the two trials in experiment 4. Parental isolate treatments shown in black and progeny treatments unshaded. Bars represent +1 S.E. Different letters above bars represent significant differences (*P* ≤ 0.05) according to a Tukey honest significant difference test. See Supplementary information table S4 for a summary of statistical analyses and Supplementary information for full details of all statistical tests.

More recently, parental isolates, C2 and C3, were shown to be homokaryon and heterokaryon (containing a population of two genetically distinct nuclei), respectively^14-16^. *In vitro*-produced homokaryon siblings cannot be genetically different from each other. Siblings of C3 are qualitatively genetically identical, containing the same alleles (Supplementary information note 5), but can display significant quantitative genetic differences in allele frequencies^15^. Variation in cassava growth might be explained by epigenetic differences among homokaryon siblings and a combination of quantitative genetic, plus epigenetic, differences among heterokaryon progeny. In order to quantify the amplitude of the effects of fungal variation among homokaryon progeny and among heterokaryon progeny on cassava productivity in the field, we conducted a fifth experiment in Kenya (one location) and Tanzania (two locations). A local cassava variety and an improved variety (for tolerance to resistance) were inoculated with five parental *R. irregularis* isolates and their single spore progeny in a randomised complete block design (Supplementary information table S3). Two of the parental isolates (C3 and A5) were heterokaryons (containing a population of two genetically distinct nuclei) and the three isolates (A1, B12 and C2) were homokaryons. Effects of inoculation treatments were not consistent across locations (Supplementary information table S6) or between two cassava varieties within a location (Supplementary information table S7). Our goal was to quantify among-AMF sibling effects on cassava growth. Thus, the analysis was separated by location, cassava variety and sibling groups. Thirty-two independent tests compared effects of a given parental isolate and its siblings on cassava root weight across all locations and with all cassava varieties (summarised in Figure 4; Supplementary figures 4-7; Supplementary information table S8). Following correction for multiple testing, 19 tests revealed a significant difference in cassava root fresh weight among plants inoculated with a given parental line and its clonal progeny. Exceptionally large differences in cassava root productivity, due to inoculation with clonally produced siblings, were observed, with up to a 3 kg plant^-1^ difference in mean root fresh weight (Figure 4). In many cases, at least one of the progeny induced significantly larger cassava root growth than the original parent. Within sites and cassava varieties, the amount of variation in cassava root productivity was similar among siblings of homokaryon parents and siblings of heterokaryon parents (Supplementary information table S9; Supplementary figure 8) indicating that quantitative genetic differences among AMF siblings are unlikely to contribute additional variation to cassava productivity. Significant differences in fungal colonization were also observed among plants inoculated with sibling fungi (Supplementary information tables S10-S11) but such effects were clearly not correlated with cassava productivity. Responses to inoculation with *R. irregularis* isolates in Africa and Colombia were highly cassava-variety dependent, showing that the genetics of the plant plays a strong role in responsiveness. This is consistent with recent studies on variation in transcriptional responses of cassava varieties to inoculation^17^. Interestingly, in this study the direction of the productivity response compared to no inoculation (known as mycorrhizal responsiveness) was extremely consistent within local cassava varieties but highly inconsistent in improved varieties (Supplementary figure 9), indicating that plant breeding may disturb cassava mycorrhizal responsiveness.

**Figure 4.**
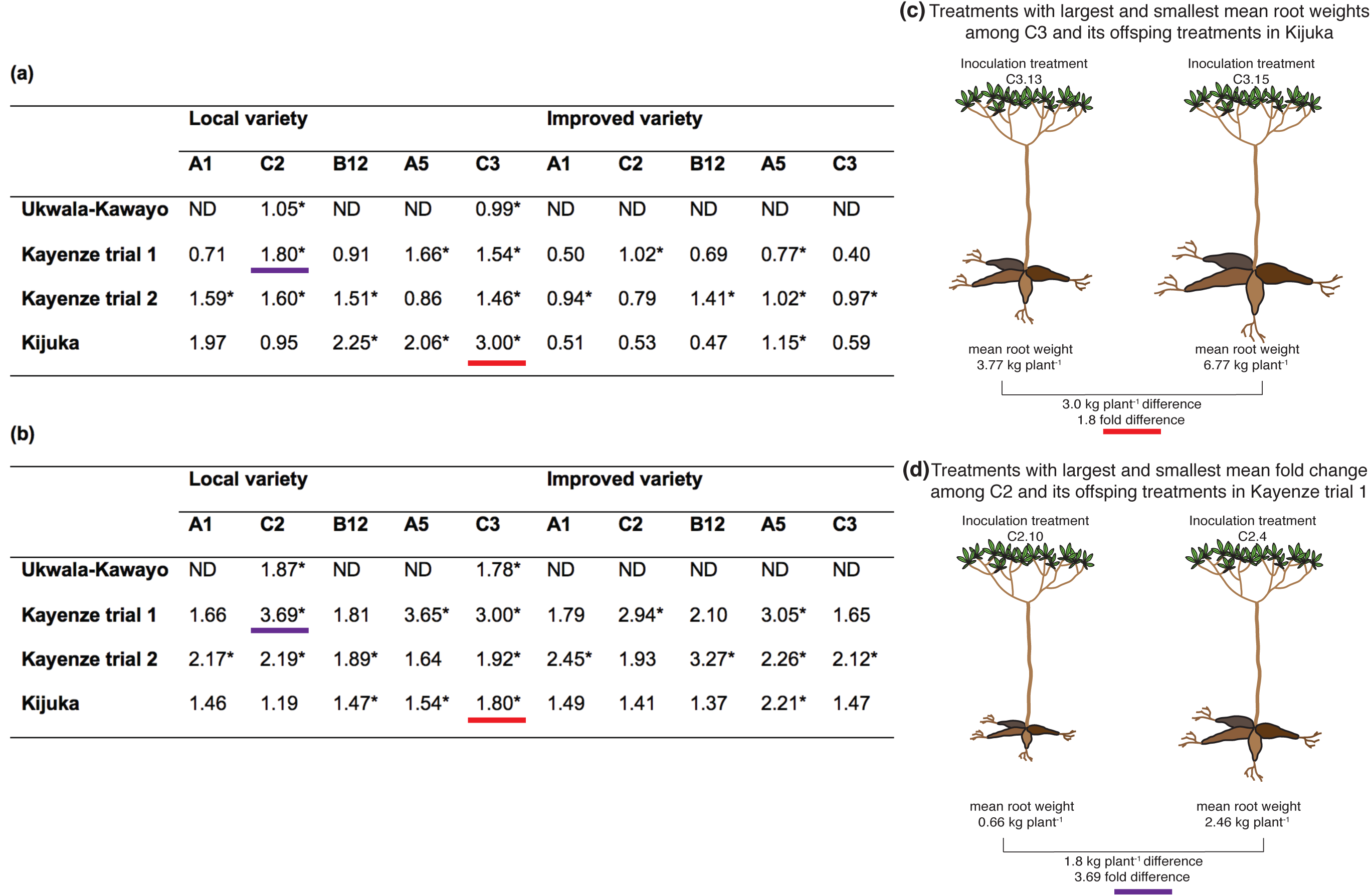
**(a)** Range of means of root fresh weight per plant (kg plant^-1^) among cassava inoculated with a given parental *R. irregularis* isolate (A1, C2, B12, A5 & C3) and its respective progeny, in four trials conducted at three locations in Kenya and Tanzania. **(b)** Fold increase of cassava root weight in the most productive treatment compared to the least productive in the same experiment. *Effect on root weight among plants inoculated with a parental AMF line and its progeny was significant at *P* ≤ 0.05. ND = no data. (c) shows the example of the treatments resulting in the largest difference in cassava root weight between two sibling progeny of the same *R. irregularis* parent underlined in red in (a) and (b). (d) shows the example of the treatments resulting in the largest fold difference in cassava root weight between two sibling progeny of the same *R. irregularis* parent underlined in purple in (a) and (b). Means for each comparison between a parental R. irregularis isolate and its progeny with a given cassava cultivar and location is provided in Supplementary figures 4-7.

This study represents several milestones. The study represents the first established link between specific patterns of genome-wide variation in a beneficial microbial species and a globally important crop species. The possibility of identifying variation responsible for increased cassava productivity, thus, opens the door to the first genetic selection program of a beneficial microorganism. A second, and more surprising milestone, is that variation among clonal offspring of an AMF parent led to additional, exceptionally large significant differences in cassava growth in real-life farming conditions in multiple locations. To put the result in context, a difference of 3kg plant^-1^ root weight would give a projected yield difference of 30 tons ha^-1^ and yet he average global yield of cassava is only 12.8 tons ha^-1^. Furthermore, the way we cultivated single-spore clonal progeny of AMF sometimes led to a more growth-promoting inoculant than the parental fungus showing that improving inoculants is a realistic possibility. There are several other reasons why these results are very unexpected (Supplementary information note 6). While The fact that cassava responsiveness to AMF is very strongly dependent on intra-specific AMF variation, the genetic identity of cassava, as well as location, highlights the extreme naivety in current use of beneficial microorganisms, such as AMF (Supplementary information note 6). We feel the current practice of using beneficial microbes to positively influence crop growth is insufficient to address the fact that use of one inoculum in one location or variety may have a positive effect, while the same “beneficial” microbe may have detrimental effects in another soil or plant variety. These data attest to the very strong potential gains of using AMF variation to improve cassava productivity. Understanding and predictably employing mycorrhizal responsiveness as a component to improve crop yield is a multi-disciplinary problem requiring expertise from microbiologists, plant breeders and agronomists alike. Future implementation of AMF in crops must consider variation in the microorganisms themselves, their effectiveness on different plant varieties and localized soil conditions and soil microbiota.

## Methods

### Fungal material used in the experiments

All *R. irregularis* isolates used in this study were originally isolated from a field in Tänikon, Switzerland. Each isolate was initiated as a single spore culture and put into *in vitro* culture with Ri-T transformed carrot roots in 2000 and subsequently maintained in the same environment and successively sub-cultured every 3-4 months in the same media^18^. The *in vitro* isolates and their origin were previously described^19^. Unless stated, all sub-culturing was conducted as in Koch et al. (2004) where colonized roots with hyphae and many spores were transferred from a 3-4 month old culture onto a Petri plate with fresh media^19^. Single spore cultures of each parental isolate were made by transferring a single spore from a 3-4 month old culture of a parental isolate onto a new plate with fresh media and a non-colonised carrot root, as described in Angelard et al. (2010)^13^. From this point on, single spore cultures were maintained by sub-culturing in the same way as parental isolates so that large amounts of clonally-grown spores could be produced from each single spore culture.

### Experiments 1 & 2: The test for a significant relationship between patterns of genetic variation in AMF and quantitative traits of AMF and plants

#### Fungal material and measurements

We cultivated *R. irregularis* isolates (A4, A5, B3, B4, B10, B15, C1, C2, C4, D1 & D4) *in vitro* in split plates and characterised fungal growth traits^13^. In experiment 1, there were 5 replicate petri plates of each isolate. We measured spore production of the isolates after 6 months of growth. To do this, we took photographs of 6 areas of 2cm^2^ in the fungal compartment of each plate using a Leica stereoscope (MZ125). An automated measurement of spore number in a given area was then made on each image with the open source software ImageJ (https://imagej.nih.gov/ij/). In addition, we measured whether the spores were more clustered together or if they were distributed more regularly, using the R package Spatstat^20^. To do this, we measured the spatial arrangement of the spores produced each fungus by measuring the nearest distance to the spores from random points chosen within each image. We also measured the hyphae produced by counting the number of hyphae that crossed two transects of 1.44 cm length. We took 5 independent photos in each dish to make this measurement.

#### Plant material and growth conditions

We propagated the cassava (*Manihot esculenta* Crantz) variety NGA-16 (CIAT, Colombia) clonally *in vitro*. Plantlets were grown in a culture chamber (25°c, 14 hours light, 90% RH) on MS medium for 1 month. Then, we placed the seedlings in an steam sterilized (180° 25-min) soil substrate (Klassman seedling substrate:perlite 1:1). After 1 month of growth *ex vitro*, we transferred the plantlets to the final steam sterilised (120°C for 40 min 2x) substrate (Klassman substrate 4:sand:clay:perlite 4:2:1:1). At this time, the plants were kept in the greenhouse at 28°C, 16 hours light, 70% RH. We inoculated each plant with 300 spores of the *R. irregularis* isolates (A3, B4, B10, C1, C2, C3, C4, C5, D1, D4 or G1). There were 15 replicates per inoculation treatment and the plants were arranged in a randomized block design. Plants were harvested after 8 months of growth. We measured the height and dry weight of aboveground and belowground parts of the plants. Dry weight was obtained after the plants were dried at 72°c for 6 days. Root colonization by AMF was determined by the grid line intersection method after clearing roots with 10% KOH for 4 hours, acidified with HCl (1%) during 5 minutes and staining with trypan blue (0.10% in a lactic acid-glycerol solution) overnight^21^.

#### Genetic relatedness among *R. irregularis* isolates

We used published ddRAD-seq data of the *R. irregularis* isolates to estimate the genetic relatedness among isolates based on single nucleotide polymorphisms^10,11^. We called SNPs following the method described in Wyss et al. 2016^10^. We then produced a matrix of presence and absence of SNPs and calculated a genetic distance matrix using the maximum distance between each comparison (supremum norm). Then we performed a hierarchical clustering analysis to infer the genetic relationship among the *R. irregularis* isolates.

#### Statistical analysis

The fungal variables spore production and extra-radical mycelial density were analyzed with a generalized mixed model using the Poisson family as a link function and the block effect was defined as random. The spore clustering, was analyzed using a non-parametric Kruskal-Wallis test. Fungal colonization was analyzed with a generalized mixed model using the binomial family as link function, we defined the block effect as random. We used a mixed linear model to analyze the plant variables: plant height, shoot dry-weight, root dry-weight and total dry-weight. The block effect was defined as a random factor in the model.

#### Phylogenetic signal

We used the R package ‘phylosignal’ in order to test whether there was a significant phylogenetic signal in fungal and plant quantitative traits and genome-wide genetic variation in the fungi^9^. We calculated two different phylogenetic signal metrics; Abouheif’s C mean and Moran’s I^22,23^. These two methods are not based on an evolutionary model and use an autocorrelation approach^24^.

### Experiment 3: A test of whether genetically different *R. irregularis* isolates differentially affect cassava growth under conventional farming conditions in Colombia

By conventional farming conditions, we mean that the cassava crop was cultivated exactly as farmers would normally cultivate cassava in this region with the same planting density, fertilization, harvesting time etc. and the only difference was that some plants were inoculated with AMF. In other words, the farmer did not have to modify the crop management in any way to allow inoculation. This is the case for experiments 3, 4 and 5 described below.

The experiment was conducted in Tauramena municipality, Casanare, Colombia (72°34’23”W, 4°57’32”N at 219 metres above sea level). The climate is tropical with average temperatures of 18°C (night) to 28°C (day), with an average air humidity of 75% and total annual precipitation of 2335 mm with 172 rain days. The soil type is an Inceptisol. Cassava (cultivar CM4574; known locally as *Cubana*) was planted as stem cuttings at a density of 10000 plants ha^-1^. There were seven different inoculation treatments where cassava plants were inoculated with one of the isolates A1, A5, B12, C2, C3 or inoculated with the inoculum carrier without fungus, or not inoculated. The experiment was set up as a randomized block design, with 6 blocks and with one replicate of each treatment per block. Each treatment in each block, was represented by nine cassava plants arranged in a plot (3 × 3 plants). Each set of 9 plants of one treatment were surrounded by a row of uninoculated plants so that any two plants at the edge of the plots of different inoculation treatments were separated from each other by two non-inoculated plants. The fungal inoculum was upscaled in an *in vitro* culture system by Symbiom s.r.o. (Lanskroun, Czech Republic) and mixed with a carrier (calcified diatomite). Stems of cassava were inoculated with 1g of the carrier, containing 1000 *R. irregularis* spores, which was placed around the stem of the cassava at planting. Plants were fertilized in total with 100 Kg ha^-1^ urea, 100 Kg ha^-1^ diammonium phosphates (DAP), 106 Kg ha^-1^ potassium chloride (KCl). Fifty percent of this was applied at 43 days after planting date and the other 50% at 61 days after planting date. There was no artificial irrigation and conventional crop management for the region was applied depending on pests, diseases and weed incidence. The bulbous roots were harvested 321 days after planting after planting and the root weight fresh weight per plant was measured. Values of the nine plants per treatment per plot were pooled and the mean value per plot was used for further statistical analysis. We tested for a significant difference in cassava root fresh weight among treatments by performing an analysis of variance using the JMP® statistical discovery software (Statistical Analysis Systems Institute, version 13). Comparison of least square means differences was done using Student’s t test at 90% of significance.

### Experiment 4: A test of whether clonally-produced single spore siblings of *R. irregularis* isolates C2 and C3 differentially affect cassava growth under conventional farming conditions in Colombia

#### Experimental design

The field experiments were established at the Utopia campus of La Salle University (72° 179 4899 W, 5° 199 3199 N) near Yopal in Casanare, Colombia with the same experiment conducted once, and then repeated in an adjacent field, in the following year. The climate is very similar to that at the location of Experiment 3. Cassava varieties used in these experiments were cultivar MCOL2737 (known locally as *Brasilera)* and cultivar CM4574. These varieties were selected because they are suitable for growing in the region and because of their growth response to AMF inoculation^8,25^. There were fourteen inoculation treatments with *R. irregularis.* This comprised the parental isolates C2 and C3 and 3 and 9 single-spore progeny of the two parental isolates, respectively. The treatments are summarized in Supplementary information table S3. Water was applied as a control in a non-inoculated treatment. Both cassava cultivars were inoculated with each of the 14 inoculation or the non-inoculated treatments in a randomized block design with nine blocks and one plant of each cultivar per treatment per block. Two rows of plants were planted around each treated plant to reduce edge effects and isolate treatments. Cassava was planted as stem cuttings (stakes) with the same methodology and planting density as in Experiment 3. There was no artificial irrigation and conventional crop management, for the region, was applied depending on pests, diseases and weed incidence. Fertilizer was applied 45 days after planting and again at 90 days after planting. Plants in the first experiment received 233 Kg ha^-1^ urea, 125 Kg ha^-1^ di-ammonium phosphates (DAP), 100 Kg ha^-1^ potassium chloride (KCl). Plants in the second experiment received 84 Kg ha^-1^ DAP, 54 Kg ha^-1^ KCl, 41 Kg ha^-1^ of Kieserite (a fertilizer comprising 3% soluble potassium, 24% magnesium and 19% sulphur) and 22 Kg ha^-1^ of Vicor**®** (a granular fertilizer comprising 15% nitrogen, 5% calcium, 3% magnesium, 2% sulphur, 0.02% boron, 0.02% copper, 0.02% manganese, 0.005% molybdenum, 2.5% zinc). Inoculation of cassava was carried out 20 days after planting. Plants were inoculated with 500 AMF propagules per plant. Each formulation with each AMF line was diluted with water to apply 10 ml to each plant close to first roots produced by the plant. Non-inoculated plants received the same volume of water. The fungal inoculum was upscaled in an *in vitro* culture system by Mycovitro S.L. (Granada, Spain) and suspended in an inert gel carrier^8^.

#### Plant and Fungal Growth Measurements

Shoot biomass was collected and weighed directly in the field (340 days after planting). Plant material was dried at 70°C until constant weight (approximately 49 hours). Cassava roots were collected and weighed in the field (360 days after planting) to measured production per plant. Root dry weight was calculated using the specific gravity method after calibration^26^. Total AMF colonization in roots was measured in fine roots with a thickness of <2 mm. Fungal structures were visualized with Schaeffer black ink^27^. The percentage of root colonization was determined by the grid line-intersect method^21^.

#### Statistical Analyses

All data were analyzed using the JMP® statistical discovery software (Statistical Analysis Systems Institute, version 10). An analysis of variance was performed to analyze data from both experiments. A mixed model was applied to test for significant differences between treatment on cassava growth or AMF colonization. In that model *Cassava variety* and *AMF line* were fixed factors and *Year* and *Block* were random factors. Restricted Maximum Likelihood (REML) was used to analyze the model with random factors. Comparison of LS Means differences was accomplished using Tukey honest significant difference or Student’s t test. To determine the variability explained by *Year* as a factor, the same model was run without this factor. AMF colonization data was *arcsin* transformed before statistical analysis

### Experiment 5: Experiment to test the amplitude of cassava root growth responses to multiple single spore progeny of *R. irregularis* parental isolates in field conditions in Kenya and Tanzania

#### Experimental design

The study was carried out in three locations: <u>Ukwala-Kawayo</u>, Siaya County, Kenya (34° 10’ 32.7” E; 00° 15’ 12.1’’ N); <u>Kayenze</u>, Biharamulo district, Tanzania (32°35’42.79”E; 02° 35’ 20.41” S) and <u>Kijuka</u>, Sengerema district, Tanzania (31° 26’ 37.18” E; 03° 12’ 3.33” S). A total of 4 trials were conducted. One in Ukwala-Kawayo, two in Kayenze, and one in Kijuka. In the trial in Ukwala-Kawayo, we used one local cassava land-race, known as Fumba Chai. In the two trials in Kayenze, we used two varieties of cassava; a local land-race known as Mzao, and an improved cultivar known as Mkombozi. In the trial in Kijuka, we used two varieties; a local land-race known as Mwanaminzi, and the improved cultivar Mkombozi. The land-races were chosen because they are grown locally by farmers. The improved cultivars were recommended by IITA as they have been bred for resistance to diseases. In Ukwala-Kawayo, we inoculated cassava with 14 different treatments with *R. irregularis*. These were: Parental isolate C2 and 5 single spore progeny. We also used isolate C5 which has been shown to be a clone of C2; Parental isolate C3 and 6 single spore progeny. In Kayenze and Kijuka, all plants were inoculated with parental *R. irregularis* isolates A1, C2, B12, A5, C3. The isolate C5 (a clone of C2) was also used. In addition, we inoculated cassava with up to 8 single-spore progeny of each of the parental isolates. All parental isolates and single spore progeny of each parental line that were used at each location and in each trial are shown in Supplementary information table S3). In addition to the plants inoculated with *R. irregularis*, there were two control treatments; no inoculation and inoculation with the carrier but with no fungus. Planting density was the same as in experiments 3 and 4. Each trial was planted as a randomized block design with treated plants surrounded by 8 non-inoculated plants as in experiment 4. There were 10 blocks in the trial in Ukwala-Kawayo and 12 blocks in all other trials. In each trial, there was one plant of each treatment combination in each block so that block also represented replicates. Methodology for planting and inoculation was the same as in experiment 4. Fertilizers were applied between 25-45 days after planting (DAP). Plants received 150 Kg.ha^-1^ N, 40 Kg.ha^-1^ P and 180 Kg.ha^-1^ K, according to the nutritional requirements proposed by the International Institute of Tropical Agriculture (IITA). Plants were inoculated at planting, as in experiment 3, with 1g of carrier diatomite containing 1000 fungal spores.

#### Plant and fungal growth measurements

At the final harvest we measured root fresh weight per plant (kg.plant^-1^) and AMF colonization in roots (% colonized root length). AMF colonization was estimated using the grid-line intersection method^21^. Additionally, in order to have a standardized measure of mycorrhizal effects on plants inoculated with different isolates, mycorrhizal responsiveness was calculated^28^.

#### Statistical analyses

Data was analysed using the R statistical software (R Core Team, 2018; version 3.5.1) and JMP® 13.2.0 (SAS institute Inc.). To test for significant differences among treatments analysis of variance (ANOVA) was performed, using a post-hoc Tukey honest significant difference (HSD) test. Where the data could not be assumed to be normally distributed we used the Wilcoxon signed-rank test.

## Supporting information

Supplemental methods, results & figures

## Data availability

The data supporting the findings of this study are available from the corresponding author upon reasonable request.

## Acknowledgements

We thank Nicolas Ruch, Jeremy Bonvin, Lucile Muneret, Dr. Tania Wyss, Dr. Christhian Fernandez, Dr. Juan Felipe Rivera and the students of the Utopia campus of Universidad de La Salle, Elias Mwangi, Robert S. Ngomuo, Joseph J. Simuda, Celvin Tibihenda, Paul Kibinza and Dr. Simon C. Jeremiah. This work was supported by the Swiss National Science Foundation (grant numbers 310030B-144079 and 31003A-162549) and by COLCIENCIAS and Swiss Confederation scholarships to IC, DCP, RP, YMO.

## Author contributions

I.C. conceived and conducted experiment 4 and analysed data; I.D.M. conceived and conducted experiments 1 and 2 and analysed the data; R.P. conceived and conducted experiment 5 and analysed the data; D.C.P-Q. conceived and conducted experiment 3 and analysed the data; C.R. conducted experiment 3, Y.M.O. conducted experiment 4; E.C.J. conducted molecular analyses on fungal cultures used as inocula; M.T. R.P. conceived and conducted experiment 5 and analysed the data; P.M.D. conducgted part of experiment 5; C.M. conceived experiment 5, B.V. conceived experiment 5, A.R. conceived experiments 3, 4 and 5, interpreted data and wrote the manuscript, IRS conceived all experiments, interpreted at the data and wrote the manuscript.

## Materials and correspondence

Correspondence or requests for materials to

## References

1 Godfray, H. C. J. et al. Food security: The challenge of feeding 9 billion people. Science 327, 812–818 (2010).

2 Rodriguez, A. & Sanders, I. R. The role of community and population ecology in applying mycorrhizal fungi for improved food security. ISME J. 9, 1053–1061 (2015).

3 Zhang, S. J., Lehmann, A., Zheng, W. S., You, Z. Y. & Rillig, M. C. Arbuscular mycorrhizal fungi increase grain yields: A meta-analysis. New Phytol. 222, 543–555 (2019).

4 Howeler, R., Lutaladio, N. & Thomas, G. Cassava: A guide to sustainable production intensification. (Food and Agriculture Organization of the United Nations, 2013).

5 Verbruggen, E. & Kiers, E. Evolutionary ecology of mycorrhizal functional diversity in agricultural systems. Evol. Appl. 3, 5–6 (2010).

6 Kiers, E. & Denison, R. Inclusive fitness in agriculture. Phil. Trans. Royal Soc. B 369 Special Issue, 1–12 (2014).

7 van der Heijden, M. G. A., Martin, F. M., Selosse, M. A. & Sanders, I. R. Mycorrhizal ecology and evolution: The past, the present, and the future. New Phytol. 205, 1406–1423 (2015).

8 Ceballos, I. et al. The *in vitro* mass-produced model mycorrhizal fungus, *Rhizophagus irregularis*, significantly increases yields of the globally important food security crop cassava. Plos One 8, doi:10.1371/journal.pone.0070633 (2013).

9 Keck, F., Rimet, F., Bouchez, A. & Franc, A. phylosignal: An R package to measure, test and explore phylogenetic signal. Ecol. Evol. 6, 2774–2780 (2016).

10 Wyss, T., Masclaux, F. G., Rosikiewicz, P., Pagni, M. & Sanders, I. R. Population genomics reveals that within-fungus polymorphism is common and maintained in populations of the mycorrhizal fungus *Rhizophagus irregularis*. ISME J. 10, 2514–2526 (2016).

11 Savary, R. et al. A population genomics approach shows widespread geographical distribution of cryptic genomic forms of the symbiotic fungus *Rhizophagus irregularis*. ISME J. 12, 17–30 (2018).

12 Sieverding, E. Vesicular-arbuscular mycorrhiza management in tropical agrosystems. (Deutche Gesellschaft für Technische Zusammenarbeit, 1991).

13 Angelard, C., Colard, A., Niculita-Hirzel, H., Croll, D. & Sanders, I. R. Segregation in a mycorrhizal fungus alters rice growth and symbiosis-specific gene transcription. Curr. Biol. 20, 1216–1221 (2010).

14 Ropars, J. et al. Evidence for the sexual origin of heterokaryosis in arbuscular mycorrhizal fungi. Nat. Microbiol. 1 (2016).

15 Masclaux, F. G., Wyss, T., Mateus-Gonzalez, I. D., Aletti, C. & Sanders, I. R. Variation in allele frequencies at the bg112 locus reveals unequal inheritance of nuclei in a dikaryotic isolate of the fungus *Rhizophagus irregularis*. Mycorrhiza 28, 369–377 (2018).

16 Masclaux, F. G., Wyss, T., Pagni, M., Rosikiewicz, P. & Sanders, I. R. Investigating unexplained genetic variation and its expression in the arbuscular mycorrhizal fungus *Rhizophagus irregularis*. bioRxiv, https://doi.org/10.1101/682385 (2019).

17 Mateus, I. D. et al. Dual RNA-seq reveals large-scale non-conserved genotype x genotype-specific genetic reprograming and molecular crosstalk in the mycorrhizal symbiosis. ISME J. 13, 1226–1238 (2019).

18 Becard, G. & Fortin, J. A. Early events of vesicular arbuscular mycorrhiza formation on Ri T-DNA transformed roots. New Phytol. 108, 211–218 (1988).

19 Koch, A. M. et al. High genetic variability and low local diversity in a population of arbuscular mycorrhizal fungi. Proc. Nat. Acad. Sci. USA 101, 2369–2374 (2004).

20 Baddeley, A. & Turner, R. spatsat: An R package for analyzing spatial point patterns. J. Stat. Softw. 12, 1–42 (2005).

21 Giovannetti, M. & Mosse, B. Evaluation of techniques for measuring vesicular arbuscular mycorrhizal infection in roots. New Phytol. 84, 489–500 (1980).

22 Moran, P. A. P. The interpretation of statistical maps. J. Roy. Stat. Soc. Series B 10, 243–251 (1948).

23 Abouheif, E. A method for testing the assumption of the phylogenetic independence in comparative data. Evol. Ecol. Res. 1, 895–909 (1999).

24 Münkemüller, T. et al. How to measure and test phylogenetic signal. Meths. Ecol. Evol. 3, 743–756 (2012).

25 Cadavid, F. Manual de nutricion vegetal: Una vision de los aspectos nutricionales del cultivo de la yuca (Manihot esculenta Crantz). 175 p (CIAT, 2011).

26 Toro, J. & Cañas, A. in Yuca: investigación, producción y utilización (ed CE Domínguez) 567–575 (PNUD-CIAT, 1983).

27 Vierheilig, H. & Piche, Y. A modified procedure for staining arbuscular mycorrhizal fungi in roots. Z. Pflanzen. Bodenk. 161, 601–602 (1998).

28 Gange, A. C. & Ayres, R. L. On the relation between arbuscular mycorrhizal colonization and plant ‘benefit’. Oikos 87, 615–621 (1999).

