## Supplemental methods, results & figures for "Using variation in arbuscular mycorrhizal fungi to drive the productivity of the food security crop cassava"

### Supplementary information

This document contains supplementary notes 1-5 and supplementary tables (tables S1-S11).

**Supplementary notes**

#### Note 1: Mycorrhizal fungal effects on plants, their use in agriculture and the justification for an AMF improvement program

Over the past decades, literally thousands of published studies, mostly in carefully controlled sterile soil conditions have shown that inoculation of hundreds of different plant species with arbuscular mycorrhizal fungi (AMF) can lead to significant increases in plant growth. Despite this, AMF are not used greatly in conventional agriculture. There are many reasons why mycorrhizal induced growth enhancement in sterile soil may not be realistic of whether the practice of inoculation is beneficial in agriculture. Firstly, crops naturally become colonized with a community of different AMF species already present in the soil. This means that the addition of an AMF inoculum has to induce growth effects above any beneficial effects already provided by the local AMF community[2]. Secondly, the inoculated fungus has to interact with a diverse microbial community and the interaction with the microbiome could potentially determine the plant growth effect. This is largely unexplored. Thirdly, growth effects induced by mycorrhizal fungi may be highly variable according to the species and genotype of the crop plant, the local soil conditions and the local AMF community, although this is poorly understood. In addition, many published studies demonstrate that different AMF species, and different isolates of the same AMF species, have strongly differential effects on plant growth from positive to negative[29,30].

Despite the above reasons, commercial AMF inoculum is typically marketed as a crop growth enhancing product. The fungal isolates have mostly not been subjected to rigorous replicated field trials in a variety of conditions allowing a prediction of which soils the product is likely to be effective in, or which species of crops or crop varieties show responsiveness. A meta-analysis of trials where grain crops were inoculated with AMF indicates that inoculation is likely to increase wheat yields overall but that effects in individual trials can be positive, negative or neutral.

Despite the enormous advances in crop yield afforded by plant improvement programs, we are not aware of any attempts to improve AMF, or other beneficial microbes, by using their natural variation (genetic or epigenetic). The AMF species *Rhizophagus irregularis* is considered a good candidate for an improvement program because the fungus can be mass-produced cleanly *in vitro*, genetically identical genotypes of the fungus can be found in agricultural soils in geographically distant locations and in a very wide variety of different environments meaning that it has the capacity to be used in a variety of different conditions and the genome of several isolates of this fungus have been fully or partially sequenced[8,11]. In our opinion, *in vitro* production of AMF inoculum is a necessity because conventional methods of non-sterile AMF inoculum production could potentially carry other unwanted microbes. Furthermore, this AMF species exhibits extremely large genetic variation among isolates and within species variation in this fungus leads to large differences in plant growth and phosphorus acquisition by plants[31-33].

**Note 2: Measurements of cassava growth traits**

In previous investigations by our group many different quantitative traits of cassava growth were measured[8]. However, the only growth trait that consistently responded significantly to inoculation with AMF was fresh and dry root weight. Furthermore, previous studies revealed that root weight was only significantly affected in the last months of the cassava crop cycle[8,12]. While it is particularly important for food production that cassava root weight can respond positively to inoculation with AMF, it is highly inconvenient for experimenters that no variables of cassava growth, that are typically measured by agronomists on above-ground characteristics of cassava before harvest, are predictors of cassava responsiveness to inoculation with AMF. For these reasons, in these multi-treatment field trials, we focussed on cassava root weight, rather than other variables of cassava growth. Although other variables were measured, as expected, none revealed any responsiveness to AMF inoculation. This is why we focus on root weight in this publication.

**Note 3: Projected yield and why we do not report yield**

The field experiments described in this study were set up as a set of biological experiments to examine the effects of variation in *R. irregularis* on the root productivity of different cassava varieties in normal farming conditions. In the experiments, a given number of replicate cassava plants were inoculated with a given inoculation treatment and randomly assigned a position in the field. Each individual cassava plant was surrounded by non-inoculated cassava plants. The reason for this design is to reduce, as much as possible, that an AMF treatment from one inoculated plant grows and colonises a nearby plant inoculated with a different fungus. With a standard planting density, typically used in cassava cultivation, this arrangement in the field means that individual inoculated plants are not affected by typical edge effects of increased light availability and less root competition that typically occurs at the edge of plots of one treatment. However, each inoculated plant was competing with the surrounding non-inoculated cassava plants, with which they could have potentially had a competitive advantage or disadvantage. Therefore, we report root productivity per plant. We are cautious not to extrapolate the productivity per plant to a yield. However, given the extremely large differences in cassava root weight per plant among different AMF parental or sibling *R. irregularis* reported in these field experiments, predicted yield differences among treatments would be extremely large. For example, in the 3$^{rd}$ experiment a predicted yield difference of up to 18 tons ha$^{-1}$ occurred between the most effective and least effective AMF isolate treatments. In the experiments in Tanzania, predicted yield differences of up to 30 tons ha$^{-1}$ occurred among cassava inoculated with fungus C3 and its offspring. We define predicted yield as the root fresh weight yield in tons ha$^{-1}$ based on the mean value of cassava root weight in a given treatment multiplied by the planting density. Obviously, because there is inevitably plant mortality in the field, this predicted yield will never be realised. However, in these experiments, we were interested in how much variation in the AMF *R. irregularis* influenced cassava root growth. Given that there were no indications that AMF inoculation treatment influenced mortality, we expect the relative differences in predicted yield among treatments to be realistic although the total yields will be smaller than predicted. In this case, predicted yields among treatments were extremely large.

**Note 4: Sources of variation in AMF affects on cassava root productivity, AMF culturing and the removal environmental effects**

In these experiments, all *R. irregularis* were cultivated *in vitro*. This allows the production of sterile fungal material of a defined isolate or progeny line of an isolate. The parental cultures (isolates A1, C2, B12, A5 and C3) were first put into *in vitro* culture in 2000 at the University of Lausanne, where they have since remained and have received identical culturing conditions. All subsequent cultivation of these isolates or their progeny single spore lines has been conducted in exactly the same *in vitro* cultivation system. We, therefore, consider that all possible environmental effects have been reduced as much as possible or eliminated. We, therefore, consider that differences in the fungal phenotype of their effects on plant growth, among isolates must be due to genetic or epigenetic sources of variation or due to somatic mutations or genomic re-arrangements by transposable elements or a combination of these factors. An experiment of genotyping using double-digest RAD sequencing (ddRAD-seq) on DNA of a parental isolate at the 1$^{st}$, 3$^{rd}$ and 8$^{th}$ generation of sub-culturing revealed only one convincing somatic mutation, indicating the genetic stability of the isolates (P. Rosikiewicz, data unpublished). Variation due to re-arrangements by transposable elements has not been investigated. In the case of clonal single spore offspring of a given parent isolate, if the isolate is a homokaryon then no genetic differences should occur among progeny, meaning that any recorded variation in plant growth could be due to epigenetic differences. While we have not recorded differences in DNA methylation marks in the genome of the single spore progeny, other work in our group indicates that DNA methylation patterns among clones of these fungi could be different (data to be published in a separate manuscript).

**Note 5**: **Single spore cultures are clonal progeny of the parent**

All sibling single spore lines of a given parental *R. irregularis* isolate used in this experiment are very likely clonally produced. This should not be confused with the debate about whether the fungus *R. irregularis* is sexual or asexual. When referring to clonal progeny we are referring to the fact that the fungi have been cultured *in vitro* in a way that allows no, or very limited possibilities, for DNA exchange or recombination. This does not mean that the fungus is biologically incapable reproducing sexually. The single spore progeny of parental *R. irregularis* isolates generated for these field trials are highly likely clonal progeny of their parents. Homokaryon isolates A1, C2 and B12 produce spores in the *in vitro* culture system that do not have the possibility to exchange DNA with any other AMF and, by definition, a homokaryon contains identical nuclei. This is supported by extensive double-digest RAD sequencing conducted in our group (data not published). In the case of single spore progeny from heterokaryon parental fungi harbouring two genetically distinct nucleus genotypes (isolates C3 and A5), the only possibility for single spore progeny to differ from their parents is that: 1. Recombination occurs between the population of two different nucleus genotypes leading to potential small qualitative genetic differences among siblings; 2. That single spore siblings of the parent do not inherit exactly the same proportion of the two nucleus genotypes from the parent, leading to different allele frequencies at bi-allelic sites in the genome among siblings. While some evidence for recombination has been observed in an *R. irregularis* isolate[34], subsequent segregation of alleles among offspring has not been shown. Double-digest RAD sequencing performed on some of the single spore progeny of C3 and A5 appear qualitatively identical to the parents, indicating that recombination was either absent or below the level detectable. In the isolate A4 (a clone of C3), little evidence of recombination has been observed among nuclei[34]. A study of progeny of C3, however, revealed that allele frequencies at bi-allelic sites, indeed, differ among sibling progeny of isolate C3[15]. For the above reasons, we conclude that in this study, single spore progeny of homokaryons are clones and cannot differ genetically while single spore progeny of the two heterokaryon isolates are qualitatively identical but can differ quantitatively in terms of variation in allele frequency.

**Note 6: Why the results are unexpected and further implications of the results for agriculture**

Experiments 4 and 5 show that sibling clonal progeny of an AMF parent leads to very high but statistically significant variation in cassava root weight. Other than the very large effect of different sibling AMF, we find several aspects of the results presented in experiments 4 and 5 to be highly surprising and to have considerable consequences for the potential use of AMF in agriculture. These are detailed below:

First, each parental fungus culture was originally initiated from a single spore which was then grown clonally *in vitro*. From each parental culture, new sibling cultures were made from spores produced by the parental culture. This means that any variation among siblings was either present in the original parental spore or has been generated while in culture. The fact that one single spore of a fungus could ultimately give rise to progeny with such a large amount of variation in their effects on plant growth is indeed surprising and unexpected.

Second, the inoculants were applied once to the crop just after planting, after which the plants grew for 1 year before harvest. Unlike most published greenhouse experiments with AMF, the soil had not been sterilised in advance. Thus, the soil would have already contained a diverse community of soil bacteria, soil fungi and AMF (made up of different AMF genera, species and different genotypes of the same species). Indeed, non-inoculated cassava became colonised by the local community of AMF. This means that an extremely small amount of the fungus (1g of carrier containing 1000 spores) was added to each plant and into an existing soil microbial community and gave rise to up to a 3kg plant$^{-1}$ difference in root weight. We find it extremely surprising that the addition of such a small amount of fungal material, one year before harvest can have such a strong effect on cassava root productivity given that the added fungus was being introduced into an already existing diverse microbial community. Our study does not allow us to establish whether the effect of the added fungus has a direct effect on cassava growth or an indirect effect through interaction or alteration with the existing microbial community. This should be the focus of future research.

Third, we find that the ability of such a small addition of almost identical sibling fungal inoculants to the soil to alter cassava growth so greatly to be slightly alarming. While some of the AMF siblings significantly increased cassava root production, others decreased root productivity. This perturbation of cassava root productivity (from positive to negative) among siblings highlights the naivety of the current use of AMF inocula. Most commercial inocula are composed of one isolate of an AMF species (often derived from one spore). Such inoculum is then used for inoculation of a variety of crop species, a variety of different crop cultivars and in a variety of different soils, with the assumption that it will improve crop production. Worryingly, we are frequently asked by farmers and bioinoculant producers, "Which species of AMF will be good for increasing cassava production?" Our current study highlights that variation even within one spore of one AMF species leads to an enormous range of plant growth responses from positive to negative. Furthermore, in each place we conducted our experiments, and with each cassava cultivar, which sibling fungus produced the highest cassava root productivity was different. This study demonstrates the enormous potential of variation in AMF to improve crop production, the fact that 2 sibling spores of the same fungus can alter cassava productivity by up to 3kg plant$^{-1}$ in a positive or negative manner suggest that at present recommending any of these inocula for use in cassava production would be highly irresponsible for the individual farmer. We hope that the future use and commercialisation of microbial inoculants will take this into consideration and that further research will allow us to predict which strain of fungus will be effective in a given location or with a given cassava variety

All unpublished ddRAD-seq data is available on request.

**References not already cited in the main manuscript**

29 Klironomos, J. N. Variation in plant response to native and exotic arbuscular mycorrhizal fungi. *Ecology* **84**, 2292-2301 (2003).

30 Sanders, I. R. & Rodriguez, A. Aligning molecular studies of mycorrhizal fungal diversity with ecologically important levels of diversity in ecosystems. *ISME J.* **10**, 2780-2786 (2016).

31 Munkvold, L., Kjøller, R., Vestberg, M., Rosendahl, S. & Jakobsen, I. High functional diversity within species of arbuscular mycorrhizal fungi. *New Phytol.* **164**, 357-364 (2004).

32 Koch, A. M., Croll, D. & Sanders, I. R. Genetic variability in a population of arbuscular mycorrhizal fungi causes variation in plant growth. *Ecol. Letts.* **9**, 103-110 (2006).

33 Chen, E. C. H. *et al.* High intraspecific genome diversity in the model arbuscular mycorrhizal symbiont *Rhizophagus irregularis*. *New Phytol.* **220**, 1161-1171 (2018).

34 Chen, E. C. H. *et al.* Single nucleus sequencing reveals evidence of inter-nucleus recombination in arbuscular mycorrhizal fungi. *eLife* **7** (2018).

**Table S1**. Statistical tests for tests of differences in fungal and plant traits among different *R. irregularis* inoculation treatments in experiments 1 and 2 and tests for a phylogenetic signal. A general linear model with Poisson distribution was applied to spore and extra-radical mycelium production in experiment 1. A Kusksal-Wallis test was applied to spore clustering data in experiment 1. A general linear model with a binomial distribution applied to mycorrhizal fungal colonization of roots in experiment 2. A general linear model with a Gaussian distribution was applied to the plant growth traits. All general linear models were mixed models with block as a random effect. Tests for a phylogenetic signal were Abouheif's $C_{mean}$ and Moran's I.

| **Fungal traits** | **Chi-squared ($\chi^2$)** | **df** | ***P*** | **$C_{mean}$** | ***P*** | **I** | ***P*** |
|---|---|---|---|---|---|---|---|
| **Spore production** | 41141 | 10 | < 0.001 | 0.377 | 0.0167 | 0.450 | 0.003 |
| **Extra-radical mycelium production** | 1114 | 10 | < 0.001 | 0.123 | 0.1122 | 0.127 | 0.082 |
| | **Kruskal-Wallis H** | | | | | | |
| **Spore clustering** | 128.54 | 10 | < 0.001 | 0.393 | 0.0115 | 0.440 | 0.004 |

| **Plant traits or fungal traits measured in the plant root** | **Res. Deviation (df 126)** | **Null res. Deviation (df 136)** | **df model - null** | ***P*** | **$C_{mean}$** | ***P*** | **I** | ***P*** |
|---|---|---|---|---|---|---|---|---|
| **Root colonization by AM fungi** | 560 | 829 | 10 | <0.0001 | 0.269 | 0.056 | 0.260 | 0.056 |
| | **F ratio $_{(10, 139)}$** | ***P*** | | | | | | |
| **Plant height** | 2.52 | 0.0082 | | | 0.272 | 0.049 | 0.121 | 0.115 |
| **Above ground plant dry weight** | 2.27 | 0.0174 | | | 0.272 | 0.046 | 0.118 | 0.132 |
| **Below ground plant dry weight** | 3.40 | 0.0005 | | | 0.526 | 0.003 | 0.370 | 0.019 |
| **Total plant dry weight** | 3.58 | 0.0003 | | | 0.534 | 0.001 | 0.375 | 0.015 |

**Table S2.** Analysis of variance performed on cassava root fresh weight in experiment 3. The data were first tested for a block effect. Given the lack of any block effect, the effect of inoculation treatment was then tested in a second one-way ANOVA with inoculation as the factor. Finally, the ANOVA was repeated with on the *R. irregularis* inoculation treatments (without the carrier or non-inoculated treatments) to confirm that significant differences in cassava root fresh weight occurred among plants inoculated with genetically different fungi.

**With data from all 7 treatments**

| **Source** | **df** | **F ratio** | ***P*** |
|---|---|---|---|
| Block | 5<br>36 | 1.39 | 0.252 |
| Inoculation treatment | 6 | 1.95 | 0.099 |
| Residual | 35 | | |

**With data from the 5 R. irregularis inoculation treatments only**

| **Source** | **df** | **F ratio** | ***P*** |
|---|---|---|---|
| *R. irregularis* inoculation treatment | 4 | 3.72 | 0.016 |
| Residual | 25 | | |

**Table S3.** Names of single spore progeny of parental *R. irregularis* isolates (A1, A5, B12, C2, C3) used in experiment 4 and in the four trials of experiment 5. Codenames of single spore progeny start with the name of the parental isolate and the number following the point denotes the single spore used to initiate the culture. Isolate C5 was also used in experiment 5. Isolate C5 has been shown to be a clone of C2 (Wyss et al.). Cassava was inoculated with the parental isolate, plus its single spore offspring in each experiment and each trial.

| | Parental isolate | | | | |
|---|---|---|---|---|---|
| | **A1** | **A5** | **B12** | **C2** | **C3** |
| **Experiment 4** | | | | C2.1 | C3.1 |
| | | | | C2.2 | C3.2 |
| | | | | C2.3 | C3.3 |
| | | | | | C3.4 |
| | | | | | C3.5 |
| | | | | | C3.6 |
| | | | | | C3.7 |
| | | | | | C3.8 |
| | | | | | C3.9 |
| **Experiment 5 (Ukwaya, Kenya)** | | | | C2.4 | C3.10 |
| | | | | C2.5 | C3.11 |
| | | | | C2.6 | C3.12 |
| | | | | C2.7 | C3.14 |
| | | | | C2.8 | C3.15 |
| | | | | | C3.16 |
| **Experiment 5 (Kayenze, Tanzania trial 1)** | A1.1 | A5.1 | B12.1 | C2.4 | C3.10 |
| | A1.2 | A5.2 | B12.2 | C2.5 | C3.11 |
| | A1.4 | A5.3 | B12.3 | C2.6 | C3.12 |
| | A1.5 | A5.4 | B12.4 | C2.7 | C3.13 |
| | A1.6 | A5.5 | B12.5 | C2.8 | C3.14 |
| | A1.7 | A5.6 | B12.6 | C2.9 | C3.15 |
| | | | B12.7 | C2.10 | C3.16 |
| | | | | C2.11 | C3.17 |
| **Experiment 5 (Kayenze, Tanzania** | A1.1 | A5.1 | B12.1 | C2.4 | C3.10 |
| | A1.2 | A5.2 | B12.2 | C2.5 | C3.11 |

| | | | | | |
|---|---|---|---|---|---|
| **trial 2)** | A1.3<br>A1.4<br>A1.5<br>A1.6<br>A1.7 | A5.4<br>A5.5<br>A5.6<br>A5.7<br>A5.8 | B12.3<br>B12.4<br>B12.5<br>B12.6<br>B12.7 | C2.6<br>C2.7<br>C2.8<br>C2.9<br>C2.10<br>C2.11 | C3.12<br>C3.13<br>C3.14<br>C3.15<br>C3.16<br>C3.17 |
| **Experiment 5 (Kijiuka, Tanzania)** | A1.1<br>A1.2<br>A1.3<br>A1.4<br>A1.5<br>A1.6<br>A1.7 | A5.1<br>A5.2<br>A5.4<br>A5.5<br>A5.6<br>A5.7<br>A5.8 | B12.1<br>B12.2<br>B12.3<br>B12.4<br>B12.5<br>B12.6<br>B12.7 | C2.4<br>C2.5<br>C2.6<br>C2.7<br>C2.8<br>C2.9<br>C2.10<br>C2.11 | C3.10<br>C3.11<br>C3.12<br>C3.13<br>C3.14<br>C3.15<br>C3.16<br>C3.17 |

**Table S4.** Results of analysis of variance using a mixed model with block and time as random effects and inoculation treatment, cassava cultivar and the interaction inoculation treatment x cassava cultivar as fixed effects in experiment 4. **a.** The analysis conducted with all inoculation treatments, including the non-inoculated control. **b.** Results of the same analysis with the non-inoculated control removed to test whether the fixed effects were significant among treatments inoculated with the fungus. Full statistical tables given in Supplementary information.

**a.**

| Variable | Variance component for time as random factor | Fixed effect source: Inoculation treatment | Fixed effect source: Cassava cultivar | Fixed effect source: Inoculation x cassava cultivar |
|---|---|---|---|---|
| Root fresh weight | 0.35% | 2.58 ($P$ = 0.0015) | 44.89 ($P \leq$ 0.0001) | 2.81 ($P$ = 0.0005) |
| Root dry weight | 0.42% | 2.74 ($P$ = 0.0007) | 93.22 ($P \leq$ 0.0001) | 3.43 ($P \leq$ 0.0001) |
| Root colonisation by AM fungi | 7.14% | 1.11 (ns) | 0.49 (ns) | 1.75 (ns) |
| Above ground fresh weight* | - | 1.01 (ns) | 5.59 ($P$ = 0.0192) | 1.94 ($P$ = 0.0254) |
| Inoculation benefit (based on root fresh weight) | 0.90% | 1.90 ($P$ = 0.0293) | 12.27 ($P$ = 0.0005) | 2.85 ($P$ = 0.0007) |
| Inoculation benefit (based on root dry weight) | 0.27% | 1.97 ($P$ = 0.0228) | 13.92 ($P$ = 0.0002) | 2.22 ($P$ = 0.0086) |

**b.**

| Variable | Variance component for time as random factor | Fixed effect source: Inoculation treatment | Fixed effect source: Cassava cultivar | Fixed effect source: Inoculation x cassava cultivar |
|---|---|---|---|---|
| Root fresh weight | 0.44% | 2.68 ($P$ = 0.0013) | 37.26 ($P \leq$ 0.0001) | 2.90 ($P$ = 0.0005) |
| Root dry weight | 0.37% | 2.89 ($P$ = 0.0005) | 81.08 ($P \leq$ 0.0001) | 3.67 ($P \leq$ 0.0001) |
| Root colonisation by AM fungi | 7.14% | 1.11 (ns) | 0.49 (ns) | 1.76 (ns) |
| Above ground fresh weight* | - | 1.06 (ns) | 5.94 ($P$ = 0.0159) | 2.05 ($P$ = 0.0203) |
| Inoculation benefit (based on root fresh weight) | 0.89% | 1.90 ($P$ = 0.0293) | 12.27 ($P$ = 0.0005) | 2.85 ($P$ = 0.0007) |
| Inoculation benefit (based on root dry weight) | 0.22% | 1.97 ($P$ = 0.0228) | 13.92 ($P$ = 0.0002) | 2.22 ($P$ = 0.0086) |

* Only measured in the $2^{nd}$ trial

**Table S5.** Summary of effects of inoculation of cassava with a parental *R. irregularis* isolate and its respective progeny (siblings). Analysis was performed separately on each cassava cultivar because strong inoculation treatment x cassava cultivar interactions occurred for nearly all variables showing that the two cultivars responded very differently to the parents and AMF progeny. **a.** Effects of C2 and its offspring. **b.** Effects of C3 and its offspring. Full statistical tables given in Supplementary information. Also see Figures 3 and Supplementary figure 3.

**a. Parental isolate C2 and its offspring**

| | MCOL2737 | | CM4574 | |
|---|---|---|---|---|
| **Variable** | **Variance component for time as random factor** | **Fixed effect source: Inoculation treatment** | **Variance component for time as random factor** | **Fixed effect source: Inoculation treatment** |
| Root fresh weight | 22.95% | 2.25 (ns) | 21.76% | 4.07 ($P$ = 0.0144) |
| Root dry weight | 19.32% | 3.06 ($P$ = 0.0365) | 23.24% | 4.76 ($P$ = 0.0070) |
| Root colonisation by AM fungi | 10.09% | 2.13 (ns) | 0.21% | 0.95 (ns) |
| Above ground fresh weight* | - | 2.74 (ns) | - | 1.24 (ns) |
| Inoculation benefit (based on root fresh weight) | 3.11% | 2.96 ($P$ = 0.0408) | 20.85% | 4.23 ($P$ = 0.0122) |
| Inoculation benefit (based on root dry weight) | 2.94% | 2.03 (ns) | 14.23% | 4.05 ($P$ = 0.0145) |

**b. Parental isolate C3 and its offspring**

| | MCOL2737 | | CM4574 | |
|---|---|---|---|---|
| **Variable** | **Variance component for time as random factor** | **Fixed effect source: Inoculation treatment** | **Variance component for time as random factor** | **Fixed effect source: Inoculation treatment** |
| Root fresh weight | 14.46% | 1.47 (ns) | 7.08% | 3.53 ($P$ = 0.0008) |
| Root dry weight | 16.65% | 1.60 (ns) | 10.99% | 3.99 ($P$ = 0.0002) |
| Root colonisation by AM fungi | 15.02% | 1.90 (ns) | 3.05% | 1.20 (ns) |
| Above ground fresh weight* | - | 1.75 (ns) | - | 1.24 (ns) |
| Inoculation benefit (based on root fresh weight) | 1.34% | 1.80 (ns) | 3.60% | 2.97 ($P$ = 0.0035) |
| Inoculation benefit (based on root dry weight) | 1.38% | 1.51 (ns) | 2.09% | 3.36 ($P$ = 0.0012) |

* Only measured in the $2^{nd}$ trial

**Table S6.** Results of a two-way ANOVA on cassava growth at final harvest in two locations, Kayenze-trial 2 and Kijuka that were planted and harvested at the same time. It considers AMF treatment and location as source of variation.

| Source of Variation | DF | Root fresh weight (kg plant$^{-1}$) | | | Main stem diameter (mm) | | | Plant height (cm) | | |
|---|---|---|---|---|---|---|---|---|---|---|
| | | SS | F | Prob > F | SS | F | Prob > F | SS | F | Prob > F |
| Treatment | 44 | 119.94 | 0.57 | 0.99 | 860.13 | 0.60 | 0.98 | 34922.43 | 0.41 | 1.00 |
| Site | 1 | 1697.67 | 357.80 | <.0001* | 4060.82 | 125.41 | <.0001* | 718485.17 | 371.05 | <.0001* |
| Treatment x Site | 44 | 116.75 | 0.56 | 0.99 | 984.36 | 0.69 | 0.94 | 30402.04 | 0.36 | 1.00 |

**Table S7.** Results of two-way ANOVA on cassava growth at the final harvest time in three trials: Kayenze-trial 1, Kayenze-trial 2 and Kijuka. It considers AMF treatment and cassava cultivar as source of variation.

| **Source of Variation** | **DF** | **Root fresh weight (kg plant$^{-1}$)** | | | **Main stem diameter (mm)** | | | **Plant height (cm)** | | |
|---|---|---|---|---|---|---|---|---|---|---|
| | | **SS** | **F** | **Prob > F** | **SS** | **F** | **Prob > F** | **SS** | **F** | **Prob > F** |
| **Kayenze trial 1** | | | | | | | | | | |
| Treatment | 42 | 33.65 | 0.64 | 0.96 | 1192.11 | 0.76 | 0.86 | 26399.78 | 0.76 | 0.87 |
| Variety | 1 | 47.61 | 47.16 | <.0001* | 861.86 | 23.22 | <.0001* | 291056.14 | 351.90 | <.0001* |
| Treatment x Variety | 42 | 39.14 | 0.74 | 0.88 | 848.67 | 0.54 | 0.99 | 36941.23 | 1.06 | 0.37 |
| **Kayenze trial 2** | | | | | | | | | | |
| Treatment | 44 | 100.49 | 1.67 | 0.0046* | 856.64 | 0.90 | 0.66 | 46777.18 | 0.68 | 0.95 |
| Variety | 1 | 181.49 | 118.62 | <.0001* | 97.28 | 4.49 | 0.0344* | 600556.03 | 383.58 | <.0001* |
| Treatment x Variety | 44 | 64.92 | 0.90 | 0.65 | 601.71 | 0.63 | 0.97 | 37395.63 | 0.54 | 0.99 |
| **Kijuka** | | | | | | | | | | |
| Treatment | 44 | 132.53 | 0.82 | 0.79 | 967.74 | 0.46 | 1.00 | 22879.69 | 0.84 | 0.75 |
| Variety | 1 | 3574.81 | 975.13 | <.0001* | 4955.18 | 102.55 | <.0001* | 373519.36 | 604.44 | <.0001* |
| Treatment x Variety | 44 | 148.75 | 0.93 | 0.61 | 718.59 | 0.34 | 1.00 | 30684.98 | 1.13 | 0.27 |

**Table S8.** ANOVA results on root fresh weight of plants inoculated by a given parental isolate and its single spore progeny at each location and with each different cassava cultivars.

| Location | Cassava Variety | Type of Variety | Test | ANOVA | | |
|---|---|---|---|---|---|---|
| | | | | F ratio | df | p-value |
| Ukwala-Kawayo | Fumba Chai | Local | C2 and its offspring | 2.1631 | 8 | 0.0463 |
| Ukwala-Kawayo | Fumba Chai | Local | C3 and its offspring | 2.0825 | 8 | 0.0475 |
| Kayenze trial 1 | Mkombozi | Improved | A1 and its offspring | 0.4578 | 8 | 0.8809 |
| Kayenze trial 1 | Mkombozi | Improved | B12 and its offspring | 0.5070 | 9 | 0.8644 |
| Kayenze trial 1 | Mkombozi | Improved | C2 and its offspring | 1.9876 | 11 | 0.0498 |
| Kayenze trial 1 | Mkombozi | Improved | A5 and its offspring | 2.1213 | 8 | 0.0469 |
| Kayenze trial 1 | Mkombozi | Improved | C3 and its offspring | 0.2621 | 10 | 0.9874 |
| Kayenze trial 1 | Mzao | Local | A1 and its offspring | 0.4272 | 8 | 0.9007 |
| Kayenze trial 1 | Mzao | Local | B12 and its offspring | 1.9678 | 9 | 0.0432 |
| Kayenze trial 1 | Mzao | Local | C2 and its offspring | 2.3612 | 11 | 0.0398 |
| Kayenze trial 1 | Mzao | Local | A5 and its offspring | 2.3854 | 8 | 0.0375 |
| Kayenze trial 1 | Mzao | Local | C3 and its offspring | 2.1689 | 10 | 0.0486 |
| Kayenze trial 2 | Mkombozi | Improved | A1 and its offspring | 2.4215 | 9 | 0.0322 |
| Kayenze trial 2 | Mkombozi | Improved | B12 and its offspring | 2.5632 | 9 | 0.0236 |
| Kayenze trial 2 | Mkombozi | Improved | C2 and its offspring | 1.2859 | 11 | 0.2413 |
| Kayenze trial 2 | Mkombozi | Improved | A5 and its offspring | 2.0236 | 9 | 0.0439 |
| Kayenze trial 2 | Mkombozi | Improved | C3 and its offspring | 2.0863 | 10 | 0.0459 |
| Kayenze trial 2 | Mzao | Local | A1 and its offspring | 2.0011 | 9 | 0.0486 |
| Kayenze trial 2 | Mzao | Local | B12 and its offspring | 2.2860 | 9 | 0.0403 |
| Kayenze trial 2 | Mzao | Local | C2 and its offspring | 1.9962 | 11 | 0.0485 |
| Kayenze trial 2 | Mzao | Local | A5 and its offspring | 0.9974 | 9 | 0.4479 |
| Kayenze trial 2 | Mzao | Local | C3 and its offspring | 2.1125 | 10 | 0.0476 |
| Kijuka | Mkombozi | Improved | A1 and its offspring | 0.3038 | 9 | 0.9717 |
| Kijuka | Mkombozi | Improved | B12 and its offspring | 0.4286 | 9 | 0.9163 |
| Kijuka | Mkombozi | Improved | C2 and its offspring | 0.3040 | 11 | 0.9837 |
| Kijuka | Mkombozi | Improved | A5 and its offspring | 2.3269 | 9 | 0.0385 |

| | | | | | | |
|---|---|---|---|---|---|---|
| Kijuka | Mkombozi | Improved | C3 and its offspring | 0.2744 | 10 | 0.9853 |
| Kijuka | Mwanaminzi | Local | A1 and its offspring | 2.0023 | 9 | 0.0443 |
| Kijuka | Mwanaminzi | Local | B12 and its offspring | 2.3952 | 9 | 0.0286 |
| Kijuka | Mwanaminzi | Local | C2 and its offspring | 0.5410 | 11 | 0.8716 |
| Kijuka | Mwanaminzi | Local | A5 and its offspring | 2.2563 | 9 | 0.0412 |
| Kijuka | Mwanaminzi | Local | C3 and its offspring | 2.4589 | 10 | 0.0236 |

**Table S9.** Tests of equal variance (four different methods) on root fresh weight of cassava inoculated with five parental *R. irregularis* isolates and their offspring in four trials conducted at three locations.

| | Local variety | | | | Improved variety | | | |
|---|---|---|---|---|---|---|---|---|
| **Test** | **DFNum** | **DFDen** | **F** | **Prob > F** | **DFNum** | **DFDen** | **F** | **Prob > F** |
| **Ukwala-Kawayo** | | | | | | | | |
| O'Brien[.5] | 4 | 191 | 0.88 | 0.47 | - | - | - | - |
| Brown-Forsythe | 4 | 191 | 0.46 | 0.76 | - | - | - | - |
| Levene | 4 | 191 | 0.51 | 0.73 | - | - | - | - |
| Bartlett | 4 | - | 0.56 | 0.69 | - | - | - | - |
| **Kayenze trial 1** | | | | | | | | |
| O'Brien[.5] | 4 | 361 | 0.23 | 0.92 | 4 | 324 | 0.57 | 0.69 |
| Brown-Forsythe | 4 | 361 | 0.63 | 0.64 | 4 | 324 | 0.77 | 0.55 |
| Levene | 4 | 361 | 1.43 | 0.22 | 4 | 324 | 1.64 | 0.16 |
| Bartlett | 4 | - | 0.59 | 0.67 | 4 | - | 1.32 | 0.26 |
| **Kayenze trial 2** | | | | | | | | |
| O'Brien[.5] | 4 | 455 | 0.85 | 0.49 | 4 | 479 | 0.93 | 0.44 |
| Brown-Forsythe | 4 | 455 | 0.60 | 0.67 | 4 | 479 | 0.38 | 0.82 |
| Levene | 4 | 455 | 1.06 | 0.38 | 4 | 479 | 0.74 | 0.56 |
| Bartlett | 4 | - | 2.43 | 0.05 | 4 | - | 1.97 | 0.10 |
| **Kijuka** | | | | | | | | |
| O'Brien[.5] | 4 | 489 | 0.47 | 0.76 | 4 | 428 | 0.32 | 0.86 |
| Brown-Forsythe | 4 | 489 | 0.74 | 0.56 | 4 | 428 | 0.12 | 0.97 |
| Levene | 4 | 489 | 0.77 | 0.55 | 4 | 428 | 0.14 | 0.97 |
| Bartlett | 4 | - | 0.47 | 0.76 | 4 | - | 0.53 | 0.72 |

**Table S10.** Range of mean colonization of cassava roots inoculated with a given parental *R. irregularis* isolate (A1, C2, B12, A5 & C3) and its progeny, in four trials conducted at three locations in Kenya and Tanzania measured at harvest.

| | **Local variety** | | | | | **Improved variety** | | | | |
|---|---|---|---|---|---|---|---|---|---|---|
| | **A1** | **C2** | **B12** | **A5** | **C3** | **A1** | **C2** | **B12** | **A5** | **C3** |
| **Ukwala-Kawayo** | - | - | - | - | 20.83* | - | - | - | - | - |
| **Kayenze trial 1** | 35.56 | 28.89 | 38.89 | 40.00* | 46.67* | 37.78 | 17.78 | 46.68* | 38.56 | 43.33* |
| **Kayenze trial 2** | 38.89 | 55.56* | 45.56* | 63.33* | 50.00* | 45.56 | 61.11* | 38.89 | 37.78 | 41.11* |
| **Kijuka** | 74.56* | 37.78* | 31.11 | 62.22* | 51.11* | 28.89* | 38.89 | 58.89* | 30.56* | 32.22 |

*Effect among plants inoculated with parental AMF line and its progeny was significant at $P \leq 0.05$.

**Table S11.** ANOVA results on AMF colonization of rootd of cassava inoculated by a given parental isolate and its single spore progeny at each location and with each different cassava cultivars.

| Location | Cassava Variety | Type of Variety | Test | ANOVA | | |
|---|---|---|---|---|---|---|
| | | | | F ratio | df | p-value |
| Ukwala-Kawayo | Fumba Chai | Local | C2 and its offspring | 0.4491 | 8 | 0.8862 |
| Ukwala-Kawayo | Fumba Chai | Local | C3 and its offspring | 2.2235 | 8 | 0.0359 |
| Kayenze trial 1 | Mkombozi | Improved | A1 and its offspring | 2.0063 | 8 | 0.0486 |
| Kayenze trial 1 | Mkombozi | Improved | B12 and its offspring | 3.2587 | 9 | 0.0125 |
| Kayenze trial 1 | Mkombozi | Improved | C2 and its offspring | 1.9857 | 11 | 0.0489 |
| Kayenze trial 1 | Mkombozi | Improved | A5 and its offspring | 3.1582 | 8 | 0.0256 |
| Kayenze trial 1 | Mkombozi | Improved | C3 and its offspring | 4.5180 | 10 | 0.0021 |
| Kayenze trial 1 | Mzao | Local | A1 and its offspring | 0.8019 | 8 | 0.6120 |
| Kayenze trial 1 | Mzao | Local | B12 and its offspring | 1.3567 | 9 | 0.2711 |
| Kayenze trial 1 | Mzao | Local | C2 and its offspring | 0.5046 | 11 | 0.8798 |
| Kayenze trial 1 | Mzao | Local | A5 and its offspring | 5.8056 | 8 | 0.0015 |
| Kayenze trial 1 | Mzao | Local | C3 and its offspring | 2.8270 | 10 | 0.0236 |
| Kayenze trial 2 | Mkombozi | Improved | A1 and its offspring | 3.7671 | 9 | 0.0078 |
| Kayenze trial 2 | Mkombozi | Improved | B12 and its offspring | 2.0983 | 9 | 0.0486 |
| Kayenze trial 2 | Mkombozi | Improved | C2 and its offspring | 2.1236 | 11 | 0.0432 |
| Kayenze trial 2 | Mkombozi | Improved | A5 and its offspring | 2.1932 | 9 | 0.0395 |
| Kayenze trial 2 | Mkombozi | Improved | C3 and its offspring | 2.0056 | 10 | 0.0479 |
| Kayenze trial 2 | Mzao | Local | A1 and its offspring | 2.1256 | 9 | 0.0421 |
| Kayenze trial 2 | Mzao | Local | B12 and its offspring | 2.9820 | 9 | 0.0223 |
| Kayenze trial 2 | Mzao | Local | C2 and its offspring | 4.3604 | 11 | 0.0020 |
| Kayenze trial 2 | Mzao | Local | A5 and its offspring | 5.2208 | 9 | 0.0020 |
| Kayenze trial 2 | Mzao | Local | C3 and its offspring | 2.9632 | 10 | 0.0211 |
| Kijuka | Mkombozi | Improved | A1 and its offspring | 2.2236 | 9 | 0.0438 |
| Kijuka | Mkombozi | Improved | B12 and its offspring | 3.2943 | 9 | 0.0142 |
| Kijuka | Mkombozi | Improved | C2 and its offspring | 1.0146 | 11 | 0.4676 |
| Kijuka | Mkombozi | Improved | A5 and its offspring | 2.0893 | 9 | 0.0489 |

| | | | | | | |
|---|---|---|---|---|---|---|
| Kijuka | Mkombozi | Improved | C3 and its offspring | 0.6706 | 10 | 0.7365 |
| Kijuka | Mwanaminzi | Local | A1 and its offspring | 4.6809 | 9 | 0.0029 |
| Kijuka | Mwanaminzi | Local | B12 and its offspring | 1.0247 | 9 | 0.4566 |
| Kijuka | Mwanaminzi | Local | C2 and its offspring | 2.1982 | 11 | 0.0403 |
| Kijuka | Mwanaminzi | Local | A5 and its offspring | 4.7106 | 9 | 0.0027 |
| Kijuka | Mwanaminzi | Local | C3 and its offspring | 2.3896 | 10 | 0.0356 |
