## Supplementary material for "Using variation in arbuscular mycorrhizal fungi to drive the productivity of the food security crop cassava": Supp fig 1

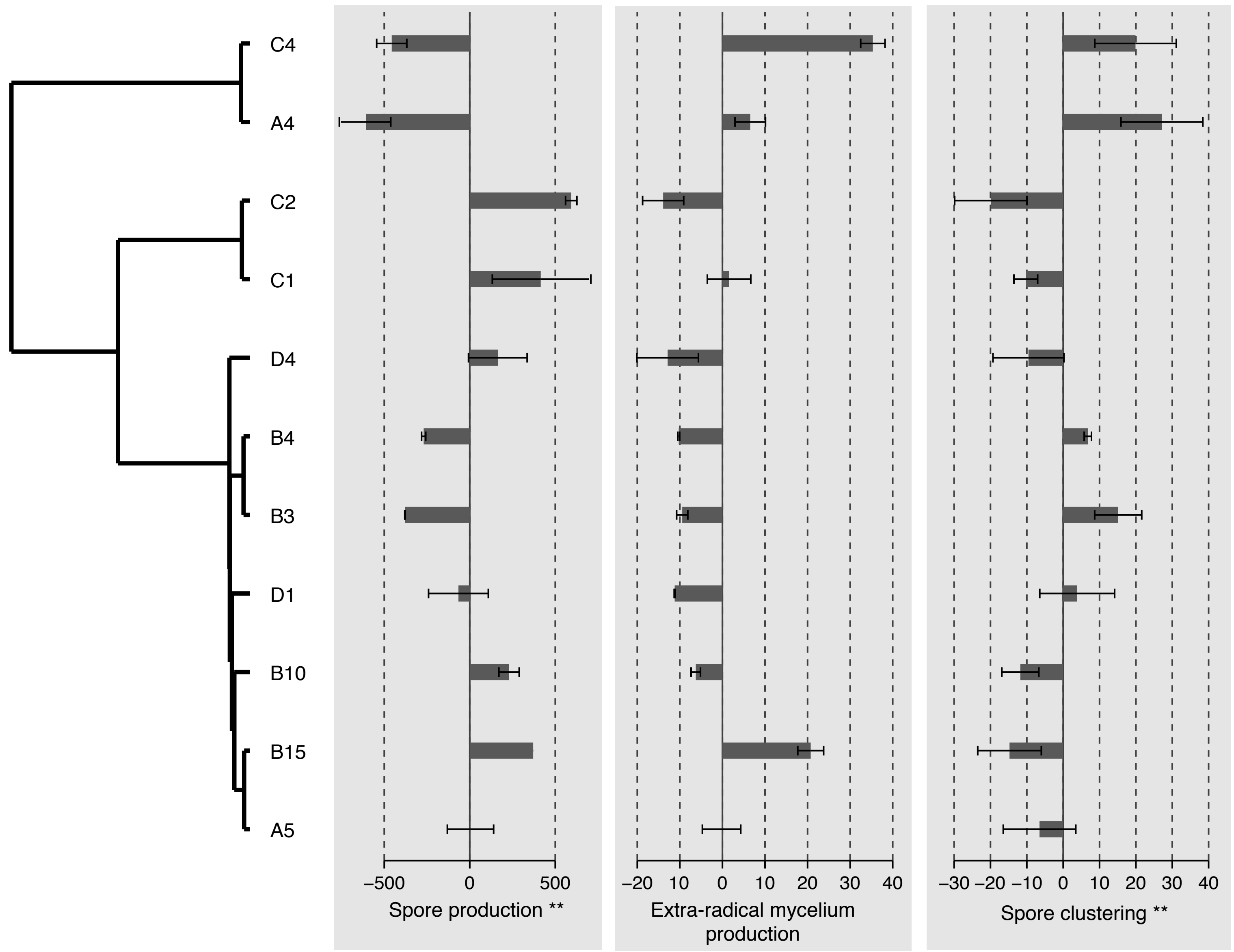


Supplementary figure 1. Variation in spore production, extra-radical mycelium production and spore clustering among 11 genetically different isolates of R. irregularis cultivated in the same in vitro environment. Values shown centred on the mean response across all treatments and bars represent the standard deviation. Value of traits of the fungal isolates are arranged according to a dendrogram of genetic relatedness among fungal isolates. * P ≤ 0.05; ** P ≤ 0.01 represent significance of tests for a phylogenetic signal of a given trait in at least one test. Statistical tests for differences in a trait among treatments and tests for the existence of a significant phylogenetic signal are given in Supplementary table S1.
