## Supplementary material for "Using variation in arbuscular mycorrhizal fungi to drive the productivity of the food security crop cassava": Supp fig 2

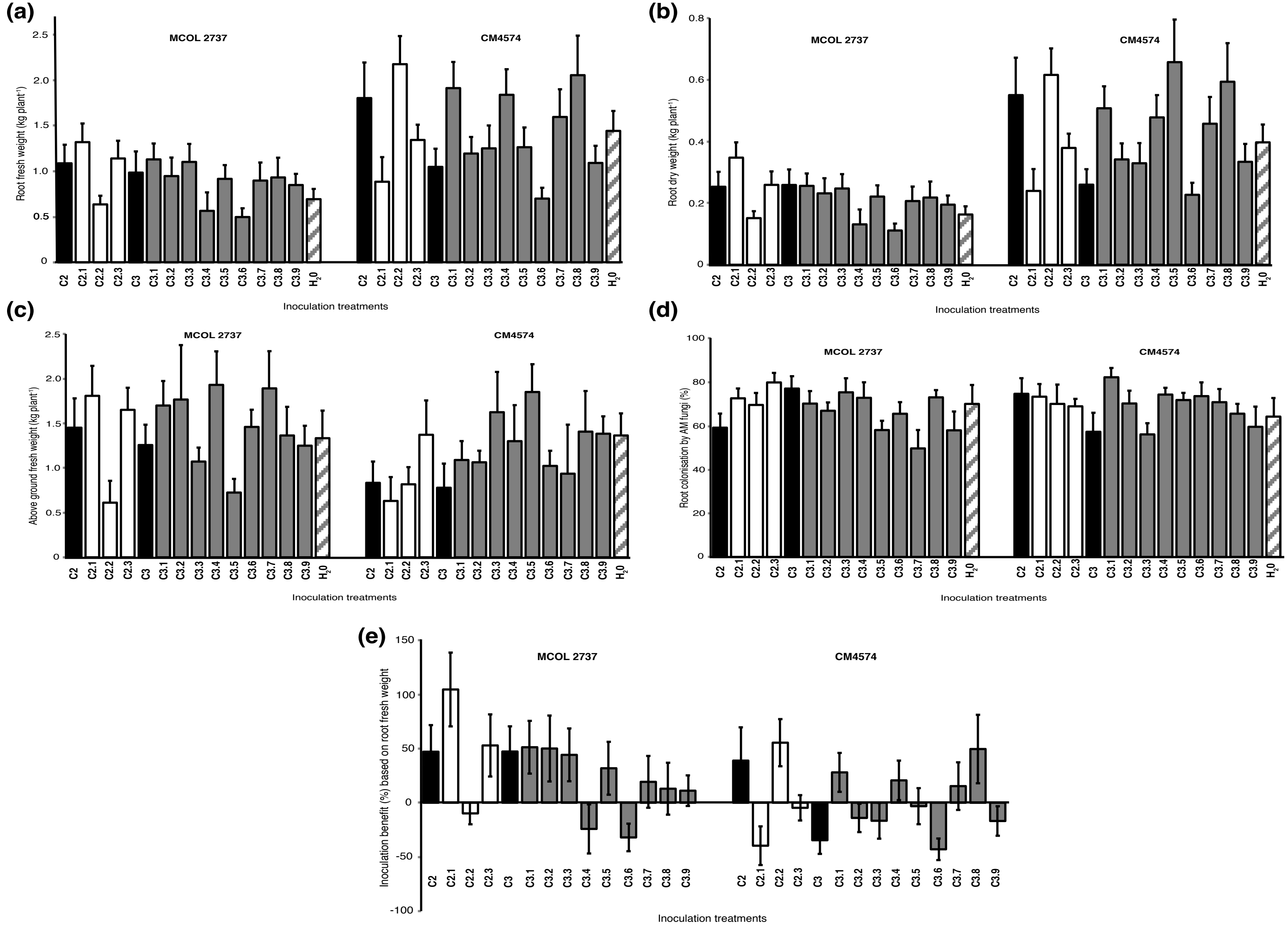


Supplementary figure 2. Effects of all inoculation treatments (including non-inoculated control "$H_20$") on cassava root fresh and dry weight, above ground fresh weight, percentage of root length colonised by AMF and inoculation benefit shown as a percentage increase or decrease of a given inoculation treatment compared to the non-inoculated control in experiment 4. All graphs represent results averaged over the two trials in experiment 4. Bars represent +1 S.E. See Supplementary table S4 for a summary of the statistical tests and Supplementary information for details of all statistical tests.
