## Supplementary material for "Using variation in arbuscular mycorrhizal fungi to drive the productivity of the food security crop cassava": Supp fig 3

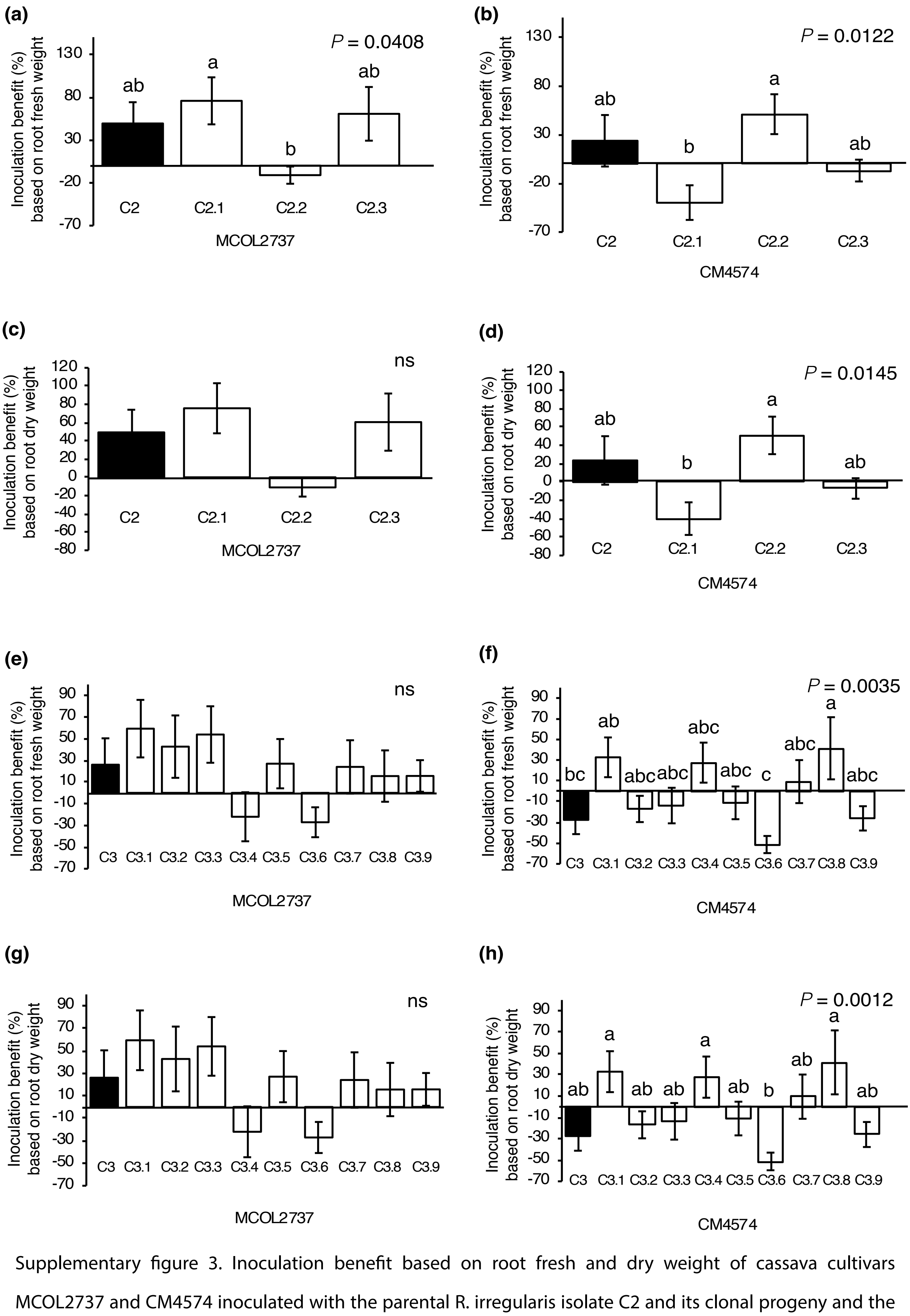


Supplementary figure 3. Inoculation benefit based on root fresh and dry weight of cassava cultivars MCOL2737 and CM4574 inoculated with the parental R. irregularis isolate C2 and its clonal progeny and the parental R. irregularis isolate C3 and its clonal progeny averaged over the two trials in experiment 4. Parental isolate treatments shown in black and progeny treatments unshaded. Bars represent +1 S.E. Different letters above bars represent significant differences ($P \leq 0.05$) according to a Tukey honest significant difference test. See Supplementary table S4 for a summary of statistical analyses and Supplementary information for full details of all statistical tests.
