## Supplementary material for "Using variation in arbuscular mycorrhizal fungi to drive the productivity of the food security crop cassava": Supp fig 4

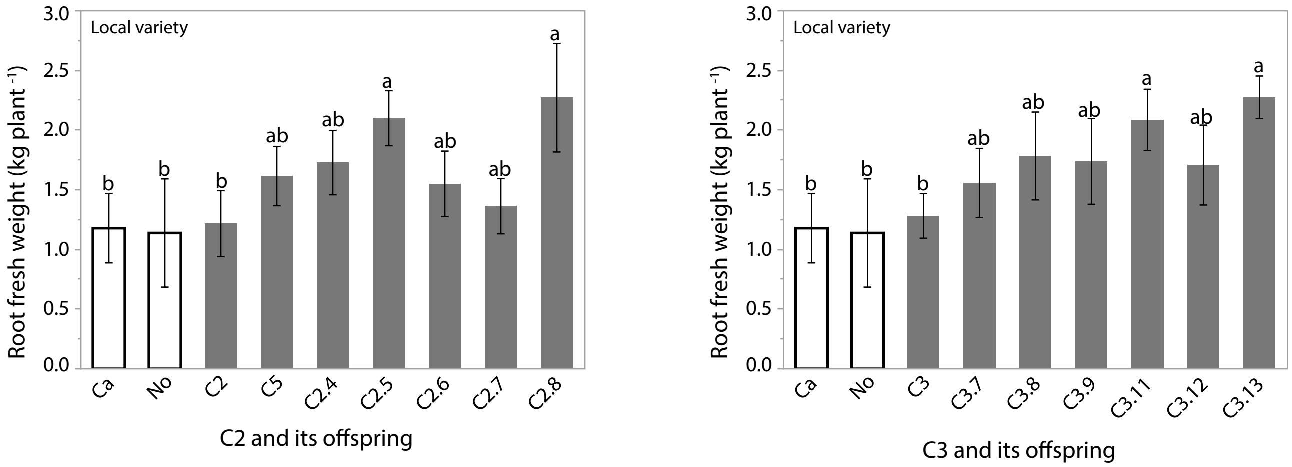


Supplementary figure 4. Root fresh weight of local cassava variety (Fumba Chai) inoculated with parental R. irregularis isolates C2, C5 and C3 and their offspring in Ukwala-Kawayo. Error bars represent $\pm$1 S.E. Different letters above bars represent significant differences at P ≤ 0.05.
