## Supplementary material for "Using variation in arbuscular mycorrhizal fungi to drive the productivity of the food security crop cassava": Supp fig 6

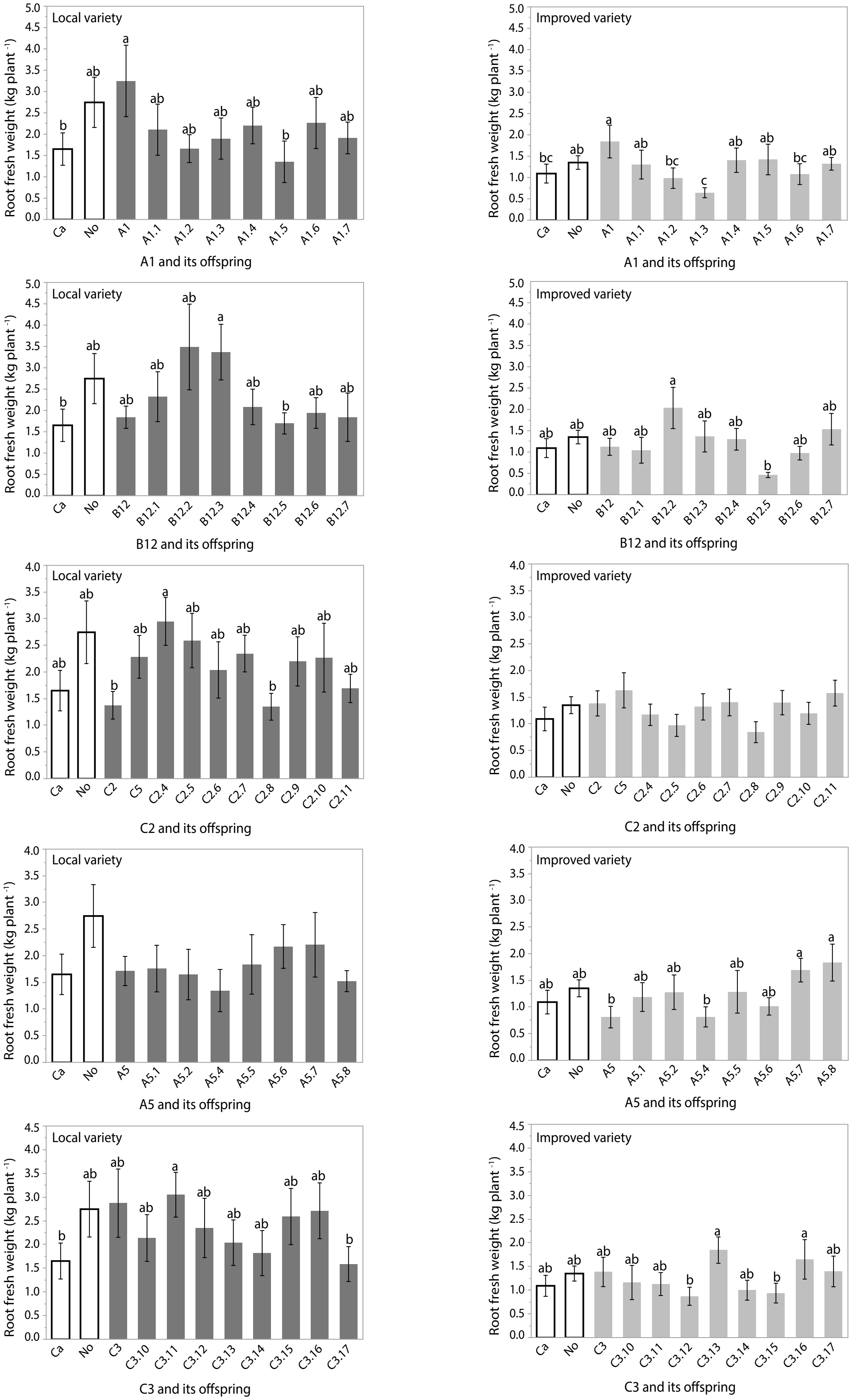


Supplementray figure 6. Root fresh weight of local cassava variety (Mzao) and improved variety (Mkombozi) inoculated with 5 parental R. irregularis isolates and their offspring in Kayenze- trial 2. Error bars represent ±1 S.E. Different letters above bars represent significant differences at $P \leq 0.05$.
