## Supplementary material for "Using variation in arbuscular mycorrhizal fungi to drive the productivity of the food security crop cassava": Supp fig 8

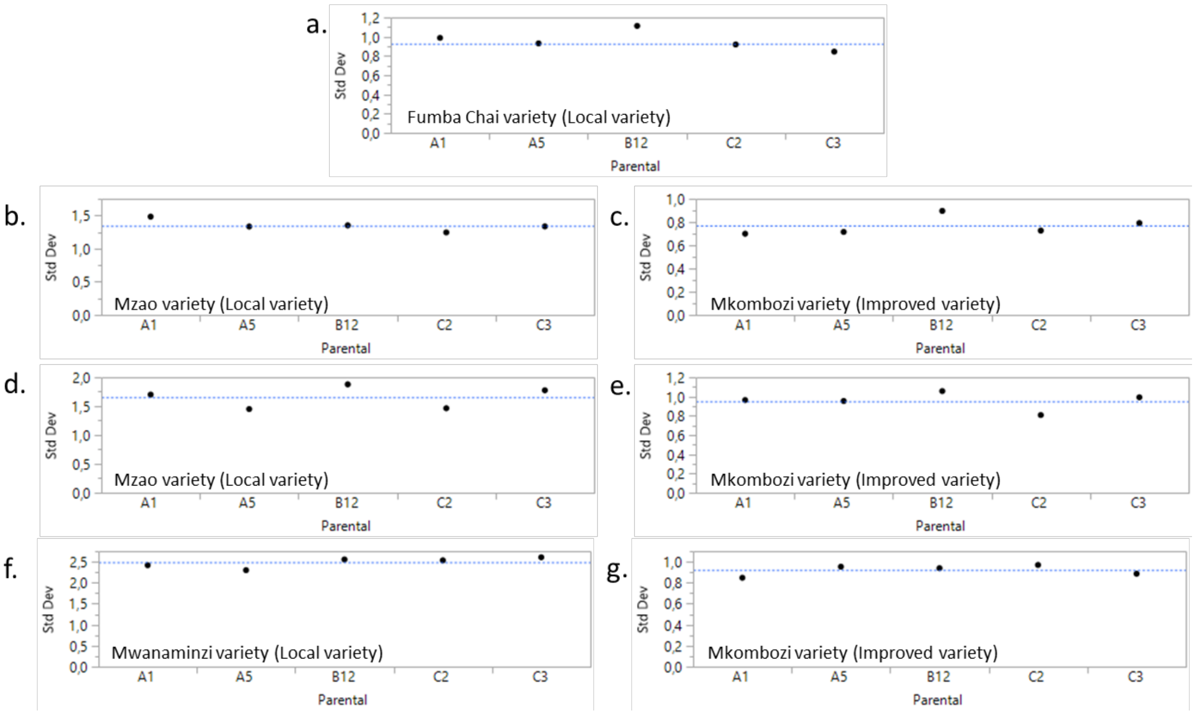


**Figure S8.** Standard deviation of root fresh weight of cassava inoculated with five parental *R. irregularis* isolates and their progeny. (a) Ukwala-Kawayo (b) Kayenze trial 1 (c) Kayenze trial 2 (d) Kijuka. Results are represented with the name of parental line: A1, A5, B12, C2 and C3. This includes the parental line itself and its progeny. The blue line represents the mean of the standard deviation.
