## Supplementary material for "Using variation in arbuscular mycorrhizal fungi to drive the productivity of the food security crop cassava": Supp fig 9

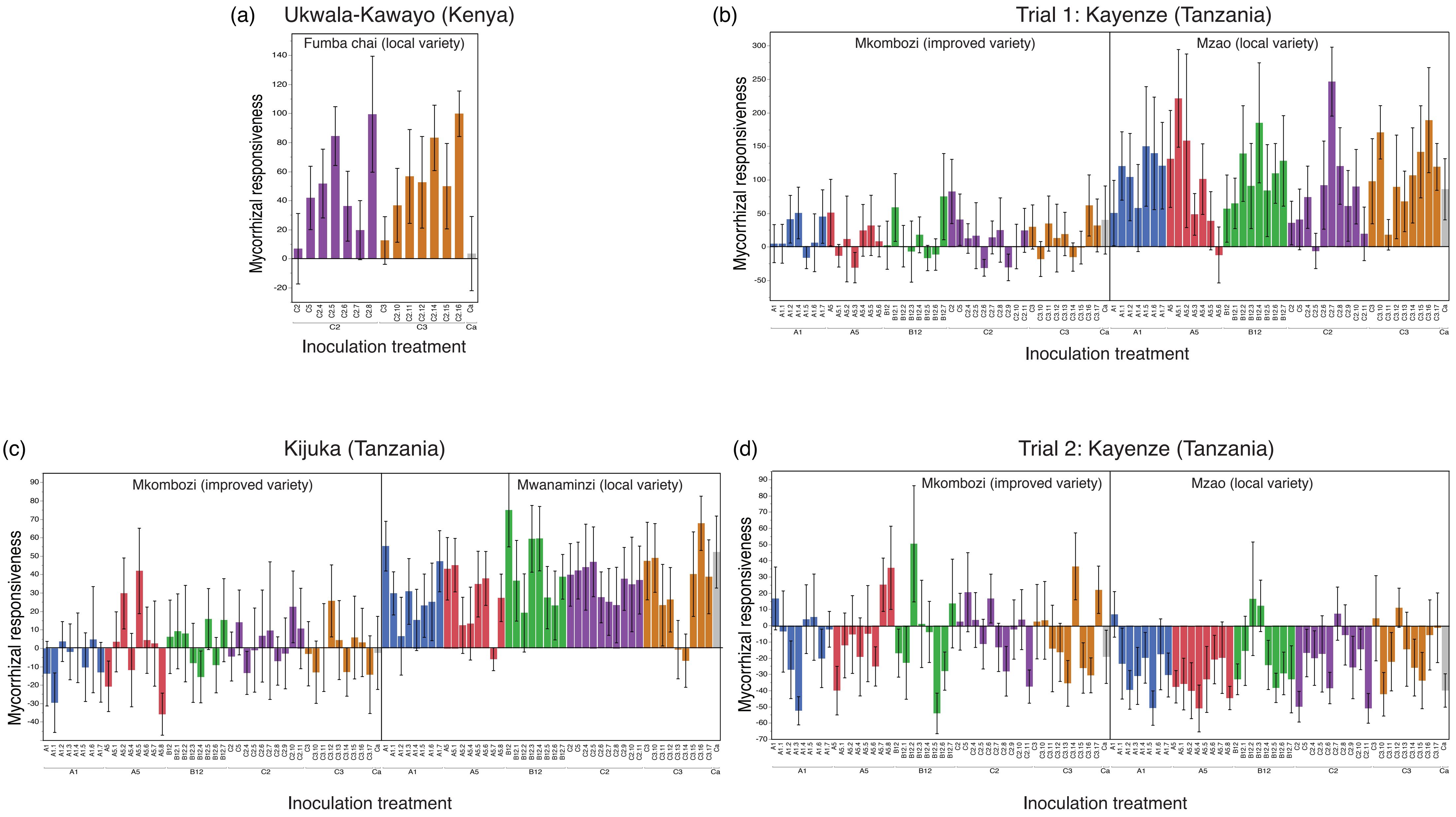


Supplementary figure 9. Mycorrhizal responsiveness of root weight of local and improved cassava varieties to inoculation with parental R. irregularis isolates and their progeny at one location in Kenya and two locations in Tanzania. The same colour bars represent inoculation with a given parental isolate and its progeny. Bars represent $\pm$1 S.E.
